# The Reelin Receptor ApoER2 is a Cargo for the Adaptor Protein Complex AP-4: Implications for Hereditary Spastic Paraplegia

**DOI:** 10.1101/2023.12.21.572896

**Authors:** Mario O. Caracci, Héctor Pizarro, Carlos Alarcón-Godoy, Luz M. Fuentealba, Pamela Farfán, Raffaella De Pace, Natacha Santibañez, Viviana A. Cavieres, Tammy P. Pástor, Juan S. Bonifacino, Gonzalo A. Mardones, María-Paz Marzolo

## Abstract

Adaptor protein complex 4 (AP-4) is a heterotetrameric complex that promotes protein export from the *trans*-Golgi network. Mutations in each of the AP-4 subunits cause a complicated form of Hereditary Spastic Paraplegia (HSP). Herein, we report that ApoER2, a receptor in the Reelin signaling pathway, is a cargo of the AP-4 complex. We identify the motif ISSF/Y within the ApoER2 cytosolic domain as necessary for interaction with the canonical signal-binding pocket of the µ4 (AP4M1) subunit of AP-4. *AP4E1*-knock-out (KO) HeLa cells and hippocampal neurons from *Ap4e1*-KO mice display increased Golgi localization of ApoER2. Furthermore, hippocampal neurons from *Ap4e1*-KO mice and *AP4M1*-KO human iPSC-derived cortical i3Neurons exhibit reduced ApoER2 protein expression. Analyses of biosynthetic transport of ApoER2 reveal differential post-Golgi trafficking of the receptor, with lower axonal distribution in KO compared to wild-type neurons, indicating a role of AP-4 and the ISSF/Y motif in the axonal localization of ApoER2. Finally, analyses of Reelin signaling in mouse hippocampal and human cortical KO neurons show that AP4 deficiency causes no changes in Reelin-dependent activation of the AKT pathway and only mild changes in Reelin-induced dendritic arborization, but reduces Reelin-induced ERK phosphorylation, CREB activation, and Golgi deployment. Altogether, this work establishes ApoER2 as a novel cargo of the AP-4 complex, suggesting that defects in the trafficking of this receptor and in the Reelin signaling pathway could contribute to the pathogenesis of HSP caused by mutations in AP-4 subunits.

## 1. INTRODUCTION

Apolipoprotein E receptor 2 (ApoER2) is a type-I transmembrane protein member of the low-density lipoprotein (LDL) receptor (LDL-R) family (Riddell, Vinogradov et al. 1999). Along with the very-low-density lipoprotein (VLDL) receptor (VLDL-R), ApoER2 functions as a membrane receptor for Reelin (D’Arcangelo, Homayouni et al. 1999, Trommsdorff, Gotthardt et al. 1999), a secreted protein that plays a central role in regulating neuronal migration and polarity during development, and neuronal plasticity in the adult brain (D’Arcangelo, Miao et al. 1995, Santana and Marzolo 2017, Wasser and Herz 2017). Reelin promotes Fyn-and Src-dependent tyrosine phosphorylation of the adaptor protein Dab1, leading to the activation of multiple signaling cascades, including PI3K-Rac1/Cdc42, PI3K-AKT-GSK3β, MEK-ERK-pCREB, and Crk/CrkL-Rap1, among others (Bock and Herz 2003, Bock, Jossin et al. 2003, Chen, Beffert et al. 2005, Lee, Chhangawala et al. 2014, Santana and Marzolo 2017). Dysfunctions in the ApoER2/Reelin signaling pathway have been documented in neurodevelopmental and neurodegenerative disorders, including Alzheimer’s disease (AD), Parkinson’s disease, Huntington’s disease, epilepsy, and autism spectrum disorder (ASD) (Fatemi 2002, Haas, Dudeck et al. 2002, Fatemi 2005, McCullough, Xu et al. 2012, Bodea, Spille et al. 2014, Baek, 2015 #635, Cuchillo-Ibanez, Mata-Balaguer et al. 2016, Cho, Kim et al. 2022). The neuroprotective role of Reelin has been recently reinforced by the discovery of a Reelin variant that delays the onset of familial AD (Lopera, Marino et al. 2023).

Reelin plays a major role in regulating neuronal polarization, migration and differentiation, all processes that rely partly on the endosomal and secretory pathways. In this context, Reelin has been shown to participate in Golgi Complex (GC) deployment within the neuronal soma towards growing dendrites (Caracci, Fuentealba et al. 2019). In Dab1-or Reelin-knock out (KO) neocortical pyramidal neurons, the GC is condensed near the nucleus, whereas in control neurons the GC extends toward the apical dendrite (Caracci, Fuentealba et al. 2019). Acute treatment of cultured hippocampal neurons and cortical slices with Reelin-conditioned medium leads to rapid deployment of the GC from the neuronal soma towards the dendrites. In turn, GC morphology and relative positioning in the neuronal soma influence axonal specification and dendritic growth as the GC tends to locate opposite the axonal domain and polarized towards the dendritic arbor (Matsuki, Matthews et al. 2010, Matsuki, Chen et al. 2013). Additionally, disruption of GC morphology directly alters neuronal migration in developing mouse neocortex and adult-born hippocampal neurons (Matsuki, Chen et al. 2013, Rao, Kirschen et al. 2018), underscoring the importance of the GC in neurodevelopment and disease.

In line with this evidence, ApoER2 participates in early brain development, and its deficiency leads to abnormal cortical development reminiscent of that in Reelin mutant mice (Reeler); nevertheless, only the loss of both ApoER2 and VLDL-R recapitulates the phenotype observed in Reeler, suggesting that both receptors participate in Reelin signaling during early development (Hack, Hellwig et al. 2007, Hirota, Kubo et al. 2018). ApoER2 interacts with NMDA receptors in the postsynaptic terminal and is necessary for the Reelin-mediated enhancement of long-term potentiation (Beffert, Weeber et al. 2005). Moreover, neuronal activity enhances the recruitment of AMPAR to the synaptic surface by recruiting an EphrinB2-ApoER2 complex (Pfennig, Foss et al. 2017). Much less is known about ApoER2 in pre-synaptic terminals, where it plays a role in calcium dynamics and neurotransmitter release (Bal, Leitz et al. 2013). Given the role of ApoER2 in both synaptic terminals, the polarized distribution of the receptor between dendritic and axonal domains is likely to be under tight regulation.

The polarized distribution of endocytic receptors of the LDL-R family depends on both their biosynthetic segregation at the *trans*-Golgi network (TGN) and post-Golgi compartments (Donoso, Cancino et al. 2009), and their post-endocytic trafficking to recycle to the correct membrane domain (Gan, McGraw et al. 2002, Donoso, Cancino et al. 2009, Sotelo, Farfan et al. 2014, Perez Bay, Schreiner et al. 2016). It would thus be expected that both biosynthetic and endocytic trafficking events influence the function and regulation of ApoER2, and the signaling efficiency triggered by Reelin binding. At the plasma membrane, ApoER2 is recognized by the monomeric adaptor protein Dab2, which binds to the NPxY motif of the receptor, leading to its internalization by clathrin-mediated endocytosis (Cuitino, Matute et al. 2005). Internalized ApoER2 is then targeted to early/sorting endosomes where it is recognized by the endosomal protein sorting nexin 17 (SNX17) (Sotelo, Farfan et al. 2014). This recognition is also dependent on the NPxY motif of the receptor and allows it to recycle back to the plasma membrane, thus avoiding lysosomal degradation. SNX17-mediated receptor recycling is important for Reelin signaling. Neurons with decreased levels of SNX17 have a reduced response to Reelin, as evidenced by a reduction in Reelin-induced dendritic outgrowth and number of dendritic spines (Sotelo, Farfan et al. 2014). Furthermore, Reelin-induced cell migration is impaired in cells with decreased levels of ApoER2 (Pasten, Cerda et al. 2015).

Although extensive information is available on the signaling and functional properties of ApoER2, as well as the characteristics of its endocytosis/degradation pathway, little is known about the receptor’s biosynthetic route, specifically its trafficking mechanisms and machinery, and the polarized distribution of ApoER2 in neurons, where it is preferentially expressed. Like other members of the LDL-R family, ApoER2 is synthesized in the endoplasmic reticulum (ER) with the assistance of several chaperones, including RAP, which is essential for its proper folding and stability (Bu and Marzolo 2000). After exiting the ER, ApoER2 traffics through the secretory pathway to reach a steady-state distribution that includes localization at the plasma membrane and compartments of the endocytic pathway. Protein transport between different compartments of the secretory and endocytic pathways is often mediated by adaptor proteins that connect the cytosolic tail of cargo proteins with the machinery that forms membrane transport carriers (Robinson and Bonifacino 2001, Bonifacino 2014). In the biosynthetic pathway, segregation into different membrane transport carriers occurs preferentially at the TGN and is based on the recognition of sorting signals in transmembrane proteins by the heterotetrameric adaptor protein (AP) complexes AP-1 and AP-4, or the monomeric protein adaptors of the GGA family (Gan, McGraw et al. 2002, Wu, Zhao et al. 2003, Gravotta, Carvajal-Gonzalez et al. 2012, Bonifacino 2014, Guardia, De Pace et al. 2018). For ApoER2, the machinery and sorting signals that mediate this segregation are unknown. However, other receptors of the family, such as LDL-R and LRP-1, are recognized by interaction of specific cytoplasmic tyrosine-based motifs (Matter, Hunziker et al. 1992, Matter, Yamamoto et al. 1994, Donoso, Cancino et al. 2009) by the µ1B subunit of the adaptor protein complex AP-1B (Ohno, Tomemori et al. 1999, Gan, McGraw et al. 2002, Donoso, Cancino et al. 2009, Guo, Mattera et al. 2013). Thus, we decided to investigate whether AP complexes recognize the cytoplasmic domain of ApoER2.

Herein, we show that the cytoplasmic domain of ApoER2 interacts with the µ4 subunit of AP-4. Mutations in any of the four subunits of AP-4 (ɛ-β4-μ4-σ4) are associated with autosomal-recessive types of Hereditary Spastic Paraplegia (HSP) referred to as AP-4– deficiency syndrome (Verkerk, Schot et al. 2009, Moreno-De-Luca, Helmers et al. 2011, Jameel, Klar et al. 2014, Abdollahpour, Alawi et al. 2015, Hardies, May et al. 2015). This congenital disorder is characterized by cerebral palsy, intellectual disabilities, microcephaly, thin corpus callosum, and seizures (Ebrahimi-Fakhari, Behne et al. 1993, Moreno-De-Luca, Helmers et al. 2011). In light of the physiologic roles of ApoER2 and its ligand Reelin in the development and function of the central nervous system (CNS), and the relatively small number of cargoes identified for AP-4 (Yap, Murate et al. 2003, Matsuda, Miura et al. 2008, Burgos, Mardones et al. 2010, Mattera, Park et al. 2017, Davies, Itzhak et al. 2018, De Pace, Skirzewski et al. 2018, Ivankovic, Drew et al. 2020, Davies, Alecu et al. 2022) (Majumder, Edmison et al. 2022), we decided to investigate this novel interaction and its physiological relevance.

## 2. MATERIAL AND METHODS

### 2.1. A detailed list of antibodies used in this study is in Supplementary Table S1

### 2.2. DNA constructs

pcDNA3-HA-ApoER2, pEGFP-HA-ApoER2, pcDNA3-RAP and Dab2-PTB were previously described (Cuitino, Matute et al. 2005). The plasmids encoding TfR-GFP and pFM4-GFP TfR were provided by Alfredo Caceres (CIMETSA and UICBC, Córdoba, Argentina). pEGFPN1-human ATG9A was described (Mattera, Park et al. 2017). The GST-fusion construct with the C-terminal domain of human µ4 (residues 160-453) was described previously (Burgos, Mardones et al. 2010).

Full-length mouse µ1A, mouse µ2, rat µ3A, rat µ3B, human µ4, as well as the C-terminal domain of human µ1B (residues 137-423) were cloned in the Y2H, Gal4-activation domain (Gal4ad) vector pACT2 (Clontech) as described (Guo, Mattera et al. 2013). The cloning of all other Y2H constructs was described previously (Aguilar et al., 2001). The generation of Y2H constructs for the expression of µ4 with single amino acid substitutions was described previously (Ross, Lin et al. 2014). To generate the µ4-GFP construct, full-length human μ4 was amplified by PCR from the previously described pCl-neo-µ4-HA (Burgos, Mardones et al. 2010, Ross, Lin et al. 2014) and cloned into the EcoRI and AgeI sites of pEGFP-N1.

To generate the Y2H ApoER2 constructs, the cytoplasmic domain of human ApoER2 was amplified by PCR using pcDNA3-HA-ApoER2 as a template and cloned in pBridge using EcoRI and SalI restriction sites.

For the generation of FM4-GFP HA-ApoER2 (Gastaldi et al., 2022), DNAs sequences from ApoER2 were amplified by PCR from previously described human HA-ApoER2 (Cuitino, Matute et al. 2005) and then cloned into the plasmid pFM4-GFP type I, which has the human growth hormone signal sequence. pFM4-GFP type I was a generous gift from Dr. Enrique Rodriguez-Boulan (Margaret Dyson Vision Research Institute, Weill Cornell Medical College. USA). Products and plasmids were digested with HindIII and Sall, and cloned into pFM4-GFP.

### 2.3. Site-directed mutagenesis

To generate replace codons for phenylalanine to alanine (F49A) in FM4-HA-ApoER2-GFP and HA-ApoER2-GFP, site-directed mutagenesis was done using the Q5® Site-Directed Mutagenesis Kit (New England Biolabs) as directed by the manufacturer. The primers were generated using Q5® Site-Directed Mutagenesis Primer design tool.

The replacement of the amino acids of the ISSF motif by alanine, for use in Y2H assays, site-directed mutagenesis was carried out on pBridge ApoER2-FL using the QuikChange mutagenesis kit (Agilent) according to the manufacturer’s instructions.

The primers used for mutations, cloning and PCR genotyping are described in supplementary **Table S2**. All the constructs were verified by DNA sequencing.

### 2.4. Yeast two-hybrid assay

*Saccharomyces cerevisiae* strain AH109 (Clontech) was maintained on yeast extract/peptone/dextrose-agar plates. Co-transformations with Gal4AD-µ4 or Gal4AD-Dab2-PH constructs and Gal4BD-ApoER2 tail constructs were done by the lithium acetate procedure using the EZ-Yeast transformation kit as described in the manufacturer’s instructions (MP Biomedicals). Transformants were plated onto yeast dropout agar medium lacking leucine and tryptophan (−2 agar plates) and allowed to grow at 30 °C, usually for 2–3 days, until colonies were visible. For colony growth assays, AH109 transformants grown on-2 agar plates were inoculated in yeast dropout liquid medium lacking leucine and tryptophan (−2 liquid medium) and allowed to grow for 16 h at 30 °C. Aliquots of the yeast liquid cultures were equalized at OD600 = 0.6 with-2 liquid medium. Drops of the equalized transformant cultures (4 µl) were deposited on top of-2 agar plates or on top of yeast dropout agar medium lacking leucine, tryptophan, and histidine (−3 agar plates) and allowed to grow at 30 °C for 3–4 days until colonies were visible. The growth of yeast colonies in-2 agar plates indicated successful co-transformation, and growth in-3 agar plates was indicative of interaction.

### 2.5. Isothermal titration calorimetry

Recombinant µ4 C-terminal domain for isothermal titration calorimetry (ITC) measurements was produced as described before (Ross, Lin et al. 2014) and dialyzed overnight at 4 °C against excess ITC buffer (50 mM Tris-HCl, 150 mM NaCl, 1 mM DTT, pH 7.4). An ApoER2 peptide (YPAAISSFDR) and an Ala-substituted peptide (YPAAASSADR) (Vivitide, MA) were also prepared in the ITC buffer. All ITC measurements were performed at 25 °C using a MicroCal PEAQ-ITC instrument (Malvern Panalytical). The chamber contained ~0.2 ml of recombinant µ4 C-terminal domain (250-500 µM), and the peptides (2.5-5.0 mM) were added in 19 injections of 4 µl each. Titration curves were analyzed using MicroCal PEAQ-ITC Analysis software (Malvern Panalytical). The binding constant and stoichiometry of the interactions were calculated by fitting the curves to a one-site model.

### 2.6. HeLa (wt and AP4E1 KO) culture and transfection

HeLa cells were cultured in high-glucose Dulbecco’s Modified Eagle Medium (DMEM) supplemented with 10% FBS and 1% penicillin/streptomycin and kept at 37°C in a 5% CO2 incubator with saturated humidity.

For immunofluorescence microscopy experiments, 15,000 cells were seeded on uncoated coverslips in a 24-well plate and grown overnight. ApoER2 plasmids were co-transfected with RAP plasmid at a 1:2 ratio in Opti-MEM medium (Gibco, Thermo Fisher) using Lipofectamine 2000 (Invitrogen). After 3-4 h of transfection, the medium was removed and replaced by fresh medium. Cells were fixed 24 h after transfection or used after 16 h for FM4 chase experiments.

For co-immunoprecipitation experiments, 1×10^6^ cells were seeded on 60mm dishes; after 24 h of growth, wt or *AP4E1*-KO HeLa cells were co-transfected with plasmids encoding µ4-GFP, RAP, and HA-ApoER2 or FLAG-ApoER2 at a 1:1:1 ratio in DMEM using polyethylenimine (PEI 25k, Polysciences). After 3-4 h of transfection, the medium was removed and replaced by fresh medium. Co-immunoprecipitation assays (see below) were performed 40 h after transfection.

### 2.7. Mouse and rat neuronal primary culture

Hippocampal and cortical neurons were obtained from C57BL/6J and C57BL/6J Ap4e1tm1b(KOMP)Wtsi mouse embryos and from Sprague-Dawley rat embryos, as previously described (Sotelo, Farfan et al. 2014, De Pace, Skirzewski et al. 2018). Animals were harvested from 17-day pregnant mice or 18-day pregnant rats. Neurons were isolated and maintained for 5-12 DIV on 35 mm coverslips (3.5×10^5^ cells/plate) coated with poly-L-lysine/laminin matrix (Sigma-Aldrich, St. Louis, MO, USA). Primary neurons were grown on Neurobasal medium (Gibco BRL, Life Technologies, USA) supplemented with B27 (Gibco, Thermo Fisher) and GlutaMAX (Gibco, Thermo Fisher). Cells were kept at 37°C in a 5% CO_2_ incubator with saturated humidity. 50,000 hippocampal neurons were seeded on poly-L-lysine/laminin-coated coverslips for transfection in 12 well plates. Transfection of ApoER2 plasmids was carried out by mixing 2 μL of Lipofectamine 2000 with plasmids encoding ApoER2 and the chaperone RAP in a 1:2 ratio in Opti-MEM medium. Culture medium was removed, supplemented with 1/3 of the original volume with full Neurobasal medium, sterile filtered and kept at 37°C. Neurons were kept in Neurobasal, and Lipofectamine plus plasmid mixed in Neurobasal medium was added dropwise to the culture plate and incubated for 1 h. The medium was then aspirated and supplemented medium was added. Half of the medium was removed every four days, and an equal volume of fresh medium was added to maintain neuronal health. Cells were fixed in paraformaldehyde (PFA) as described below after 24 h of expression unless otherwise specified.

### 2.8. iPSCs culture and differentiation to i3Neurons

Human iPSCs were cultured and differentiated to i3Neurons as previously described (Fernandopulle, Prestil et al. 2018) with some modifications. Briefly, iPSCs were cultured on Matrigel-coated (Corning) dishes in Essential 8 medium (Gibco, Thermo Fisher) with supplements, antibiotics, and ROCK inhibitor (Tocris Bioscience). When 70-80% confluent, iPSCs were passaged with Accutase and seeded on Matrigel-coated dishes in Induction Medium (DMEM/F12 HEPES with N2 supplement, NEAA (Non-Essential Amino Acids) Solution, GlutaMAX (Gibco, Thermo Fisher) and Doxycycline (DOX)). The induction medium was changed every day for three days. Then, cells were dissociated with Accutase (Gibco), counted, and seeded on poly-L-ornithine-coated dishes in Cortical Medium (BrainPhys Neuronal medium (STEMCELL Tech.) supplemented with 1x B27 (Gibco, Thermo Fisher), 10 ng/mL BDNF (PreproTech), 10 ng/mL NT-3 (PreproTech) and 1 µg/mL Laminin (Gibco, Thermo Fisher) plus DOX. Half of the medium was removed every 3-4 days or once a week, and fresh medium was added to maintain i3Neurons until experiments were performed.

### 2.9. Generation of AP4M1-knock-out cells using CRISPR–Cas9

Two sgRNAs (sgRNA1 5′-ACACGCGTTCTTTTGTTCCG and sgRNA2 5′-AGACGAGTCCCCGGTTGTCA) targeting *AP4M1* were designed using CHOPCHOP (https://chopchop.cbu.uib.no/) and cloned into pSpCas9(BB)-2A-GFP (pX-458) vector (Addgene plasmid #48138) at the BbSI restriction site as described (Ran, Hsu et al. 2013). sgRNAs were co-transfected into iPSCs using Lipofectamine 2000 (Invitrogen) and GFP positive cells were sorted 48 h later into 96-well plates. Single cells were allowed to grow in a complete E8-Flex medium (Gibco, Thermo Fisher) supplemented with RevitaCell (Gibco, Thermo Fisher) for 1-2 weeks. Single colonies were then transferred to 6-well dishes and grown to confluency. Homozygous clones were then identified by PCR genotyping, yielding a 677 bp fragment for the WT cells and ~350 bp fragment for the CRISPR-edited cells. The PCR products were run on a 1% agarose gel, and positive clones were selected and expanded.

### 2.10. Recombinant Reelin

Recombinant mouse Reelin was obtained from HEK 293 cells stably expressing the full-length protein. Cells were cultured to produce a Reelin-conditioned medium as described (Sotelo, Farfan et al. 2014). The mock-conditioned medium was prepared using the same protocol from control HEK 293 cells. Briefly, cells were cultured until 80% confluent in high-glucose DMEM with 10% FBS containing penicillin and streptomycin and 0.5 mg/mL of G418 (Gibco, Thermo Fisher) at 37°C. After washing twice with PBS, the cells were cultured in high-glucose DMEM without supplements or Neurobasal medium without phenol red (Gibco) for an additional 24 h. The cell medium was collected and centrifuged at 1,000 rpm for 5 min, and the supernatant was stored at 4°C. This procedure was repeated two more times. The collected medium was concentrated using Amicon ultra-15 centrifugal filter units (filter membrane, 100 kDa).

### 2.11. Co-immunoprecipitation

Co-immunoprecipitation was performed 40h after co-transfection. HeLa cells were washed twice with 1X cold PBS and lysed on ice for 10 min with lysis buffer (0.5% Nonidet P40, 75mM NaCl, 50mM Tris-HCl pH 7.4, 1mM EDTA with protease inhibitors) followed by centrifugation at 14,000 rpm at 4 C° for 15 min. The supernatant was separated into two fractions, 10% of the volume for input, and the remainder was incubated on a rotator with 20 μl of GFP-Trap agarose beads (CromoTek), previously washed 3 times with lysis buffer, at 4°C for 1 h. After incubation, the beads were collected by centrifugation at 2,000 rpm at 4°C and then washed three times with lysis buffer before being resuspended in 25 µl of 2X loading buffer for subsequent SDS-PAGE and immunoblot analysis.

### 2.12. Immunofluorescence microscopy

Mouse and rat primary cultured neurons, HeLa cells and fibroblasts were washed with PBS and fixed with 4% PFA in PBS for 10 min at room temperature. Cells were washed 3 times for 5 min with PBS and then permeabilized for 10 min with 0.2% v/v Triton X-100 at room temperature. Then cells were blocked for 30 min with PBS containing 0.2% gelatin. Antibodies were diluted in the same blocking reagent and incubated in a wet chamber for 30 min at 37°C. Next, coverslips were washed 3 times with 1X PBS before incubation with Alexa Fluor secondary antibody as described in the previous step. After washing with 1X PBS, coverslips were mounted with Fluoromount-G (Electron Microscopy Sciences).

I3Neurons were differentiated for 12 days (Golgi deployment experiments) or 30 days on poly-L-ornithine-coated glass coverslips (2-3 ×10^5^ cells/well). Neurons were washed with PBS, fixed with 4% PFA and 4% sucrose, and permeabilized for 10 min with 0.2% v/v Triton X-100 at room temperature. For pCREB detection, the PBS buffer was supplemented with phosphatase inhibitors (1 mM orthovanadate, 1 mM glycerol 3 phosphate, 5 mM sodium fluoride) for every step. After blocking with 5% bovine serum albumin (BSA), cells were incubated in a wet chamber at 4°C with primary antibodies overnight. The secondary antibodies, Alexa Fluor 488-, 555-or-647-conjugated, were incubated in a wet chamber at room temperature. Fluoromount-G mounting medium with DAPI (Invitrogen, Thermo Fisher) was used.

### 2.13. Microscopy and image analysis

Confocal microscopy images were collected using Zeiss LSM 780 or Zeiss LSM 880 confocal microscopes with a Plan Apochromat 63x objective (N.A. 1.40) or a 40x objective for polarity index analysis. Pearson’s co-localization coefficient was calculated using JaCOP co-localization plugin in Fiji. Sholl analysis was calculated using Simple Neurite-Tracer plugin in Fiji. Intensity profiles were obtained by drawing a 20-μm line and Plot Profile tool in Fiji. Data were then plotted using Prism 8 software. For i3Neurons, images were acquired with Nikon Eclipse Ti2 inverted microscope and Leica DM2000 microscope with Axiocam 202 mono Zeiss microscope camera.

### 2.14. Immunoblotting

DIV 12-14 cortical neurons were scraped from the plate in lysis buffer (PBS pH 7.5, 0.5% v/v Triton X-100 supplemented with protease inhibitors (EDTA-free Complete; Roche), kept on ice for 15 min and then centrifuged at 15,000 x *g* for 10 min. Samples were denatured at 95 °C for 5 min in Laemmli sample buffer (Bio-Rad) containing 2.5% v/v 2-mercaptoethanol (Sigma-Aldrich), then resolved by SDS-PAGE and transferred onto PVDF membranes (ThermoFisher). Membranes were blocked with 5% w/v non-fat milk (Bio-Rad) in Tris-buffered saline (TBS) (KD Medical) containing 0.1% v/v Tween 20 (Sigma Aldrich), probed with different primary and HRP-conjugated secondary antibodies, and imaged with SuperSignal West Dura Extended Duration Substrate (Thermo Fisher).

I3Neurons were differentiated on poly-L-ornithine-coated 6-well plates (1×10^6^ cells/well). After the indicated differentiation days, neurons were washed with PBS and lysed with lysis buffer (1x PBS 1% Triton X-100, protease and phosphatase cocktail A32959, Pierce™ Mini Tablets) then centrifuged at 15,000 x *g* for 10 min. The supernatant was collected, quantified, and prepared with a loading buffer before SDS-PAGE. Samples for ATG9A detection were incubated at 37 °C for 10 min. Other samples were incubated at 95°C for 5 min. After SDS-PAGE, gel samples were transferred onto PVDF membranes (Thermo Fisher), blocked, and immunoblotted. Images were acquired with a UVITEC system and analyzed with Fiji.

### 2.15. Biotinylation of neurons

Mouse hippocampal neurons (wt and *Ap4e1*-KO) or human i3Neurons (differentiated for 30 days), seeded on poly-L-lysine-coated 6-well plates or on poly-L-ornithine-coated 6-well plates, respectively, as described were surface biotinylated. The neurons were washed in ice-cold PBS and biotinylated with 0.2 mg/mL biotin (Thermo Fisher, #21335) at 4°C for 15 min. Then, biotin was quenched with 50 mM Tris pH 7.5 and 100 mM NaCl twice for 10 min on ice. The neurons were lysed with lysis buffer (PBS, 2% Triton-X100 plus protease inhibitors), centrifuged, and incubated with streptavidin beads (Pierce, #20349) washed in lysis buffer containing 0.3% SDS at 4°C for 2h. The beads were washed three times sequentially in PBS 1% Triton-X100, PBS 1% Triton-X100 1M NaCl, and PBS. All traces of the wash buffer were removed, and the beads were re-suspended in loading buffer. The biotinylated proteins were analyzed by SDS-PAGE and immunoblotting.

### 2.16 RNA extraction and quantitative RT-PCR

Human i3Neurons were differentiated for 30 days on poly-L-ornithine-coated 6-well plates (4×10^5^ cells/well). Total RNA was extracted using TRIzol reagent (Invitrogen) and 1 µg of RNA was reverse transcribed with RevertAid First Strand cDNA Synthesis Kit (Thermo Scientific #K1622). Real-time PCR was performed in a QuantStudio 3 Real-Time-PCR-System (Applied Biosystems, Thermo Fisher Scientific) using Maxima SYBR Green/ROX qPCR Master Mix (2X) (Thermo Scientific #K0221) with 200 nM of ApoER2 primers. The thermocycler conditions were denaturation at 95 °C for 15 s, annealing at 55 °C for 30 s and extension at 72 °C for 30 s. The expression levels of ApoER2 were normalized to GAPDH expression using the delta–delta Cq method (2−ΔΔCq). The primers used are listed in Supplementary Table S2.

### 2.17. FM4 chase experiments

HeLa cells and hippocampal neurons were transfected with plasmids encoding FM4 recombinant receptors for 15 h and then treated with 2 μM DD-Solubilizer (TakaraBio/Clontech) to induce the release of the receptors from the ER into the secretory trafficking, as previously described (Al-Bassam, Xu et al. 2012). After different times of DD-Solubilizer treatment, cells were fixed and used for immunofluorescence.

### 2.18. Polarity index measurement

The polarity index was measured as previously described (Farias et al., 2012) (Guo et al., 2016). In brief, one-pixel lines were drawn along three dendrites and a representative mid-to-distal axon segment trace starting 30 µm from the neuronal soma and ending at the axonal tip identified either by GFP or HA staining. The fluorescence intensities of at least 4 dendritic segments were averaged, and the dendrite/axon polarity index was calculated for each neuron. Axonal identity was confirmed either by the absence of MAP2 staining or the presence of ANKG staining. A polarity index of 1 represents nonpolarized distribution; a polarity index <1 represents preferential axonal localization; a polarity index >1 represents preferential dendritic localization. Analyses were made using Fiji segmented line tracer and intensity measure.

For heatmaps, full lengths of dendrites traced from the soma and full lengths of the axon drawn 30 µm from the neuronal soma and ending at the axonal tip, were identified either by GFP or HA staining for each neuron in the analysis. Gray values were measured every 0.08 μm using the Plot Profile function in Fiji. Regardless of dendrite or axonal length, gray values were averaged in 10 bins of the same size encompassing the full length of the neurite from the soma to the end of the segment. Averaged gray values were then plotted for each axon and averaged in a Heatmap using Prism.

### 2.19. Analyses of Reelin-induced effects

To detect nuclear pCREB, i3Neurons were starved of supplements for 2 h with Hanks’ medium and stimulated with 10 nM of Reelin or equivalent mock volume for 30 min at 37°C. At the end of the treatments, cells were washed in 1x PBS in the presence of phosphatase inhibitors, fixed, and immunostained with anti-phospho-CREB Ser133 (Cell Signaling Technology, 87G3 Rabbit mAB #9198) and anti-MAP2, and for nuclear staining with DAPI. Epifluorescence images were analyzed using Fiji software, the integrated intensity was measured within the nuclei (region of interest), and the background intensity was subtracted to obtain the final value of each nuclear pCREB in all the images analyzed. Values were ranked and the intensity of each nuclear pCREB was plotted.

The phosphorylation of AKT and ERK was determined by immunoblotting to evaluate the Reelin signaling pathway. The cells were depleted of supplements for 2 h and stimulated with Reelin (or mock as control) for 20 and 40 min at 37°C in BrainPhys neuronal medium for i3Neurons or Neurobasal medium for mouse primary cortical neurons. Neurons were washed in ice-cold PBS and lysed in PBS, 1% Triton X-100 containing protease and phosphatase inhibitors (A32959, Pierce™ Mini Tablets). Phosphorylation of AKT and ERK were evaluated after analysis by SDS-PAGE followed by immunoblotting. The activation rate of each pathway was calculated using data from phosphorylated and total proteins normalized to actin. The ratios from the starved samples were subtracted from the ratios corresponding to the treatments with mock or Reelin. Final numbers were plotted and analyzed with GraphPad.

To analyze Golgi deployment, DIV5 mouse hippocampal neurons were incubated with 20 nM Reelin or Mock medium for 2 h after 4 h of incubation in depletion medium (only neurobasal medium (Gibco, Thermo Fisher) plus 1% GlutaMAX). i3Neurons differentiated for 12 days were starved for 2 h and treated with 10 nM Reelin or mock-treated for 30 min. Cells were fixed and stained with anti-GM130 and DAPI. Epifluorescence images were analyzed using Fiji software. For the deployment measurements, a straight line was drawn using the straight-line tool from the outermost edge of the nucleus limiting with the GC to the outermost signal of the Golgi marker. The length of the line was measured in each condition and plotted.

### 2.20 Statistical analysis

The immunofluorescence images and immunoblots were quantified with Fiji software and detailed in the figure legends. Data were analyzed using GraphPad Prism. Values report the mean ± SEM (standard error of the mean) or SD (standard deviation). Violin plots show median (bold dashed line) and quartiles (dotted lines). Mann-Whitney’s test was used to evaluate the statistical significance between two conditions. ANOVA with Tukey’s test or Sidak’s test were used for multiple comparisons.

## 3. RESULTS

### 3.1. ApoER2 interacts through a cytosolic sequence containing an IXXF/Y motif with a canonical signal-binding site on the µ4 subunit of AP-4

To assess the role of AP complexes in the trafficking of ApoER2, we performed a yeast two-hybrid (Y2H) screen testing the interaction of the cytosolic domain of the receptor (residues 849-963) with the signal-binding µ subunits of the AP-1, AP-2, AP-3, and AP-4 complexes. As a positive control, we confirmed that the cytosolic tail of the human ApoER2 interacts with the endocytic adaptor Dab2 (Fig. 1A) (Cuitino, Matute et al. 2005). We found robust interaction of the ApoER2 tail with the µ1A and µ1B subunit isoforms of the AP-1 complex, and with the µ4 subunit of the AP-4 complex (Fig. 1 B). In addition, we detected a weak interaction with the µ2 subunit of the AP-2 complex and no interaction with the µ3A and µ3B subunit isoforms of the AP-3 complex (Fig. 1 B). Due to the importance of both ApoER2 and AP-4 in the development and function of the CNS, we decided to study the ApoER2–AP-4 interaction in more detail.

**Figure 1.**
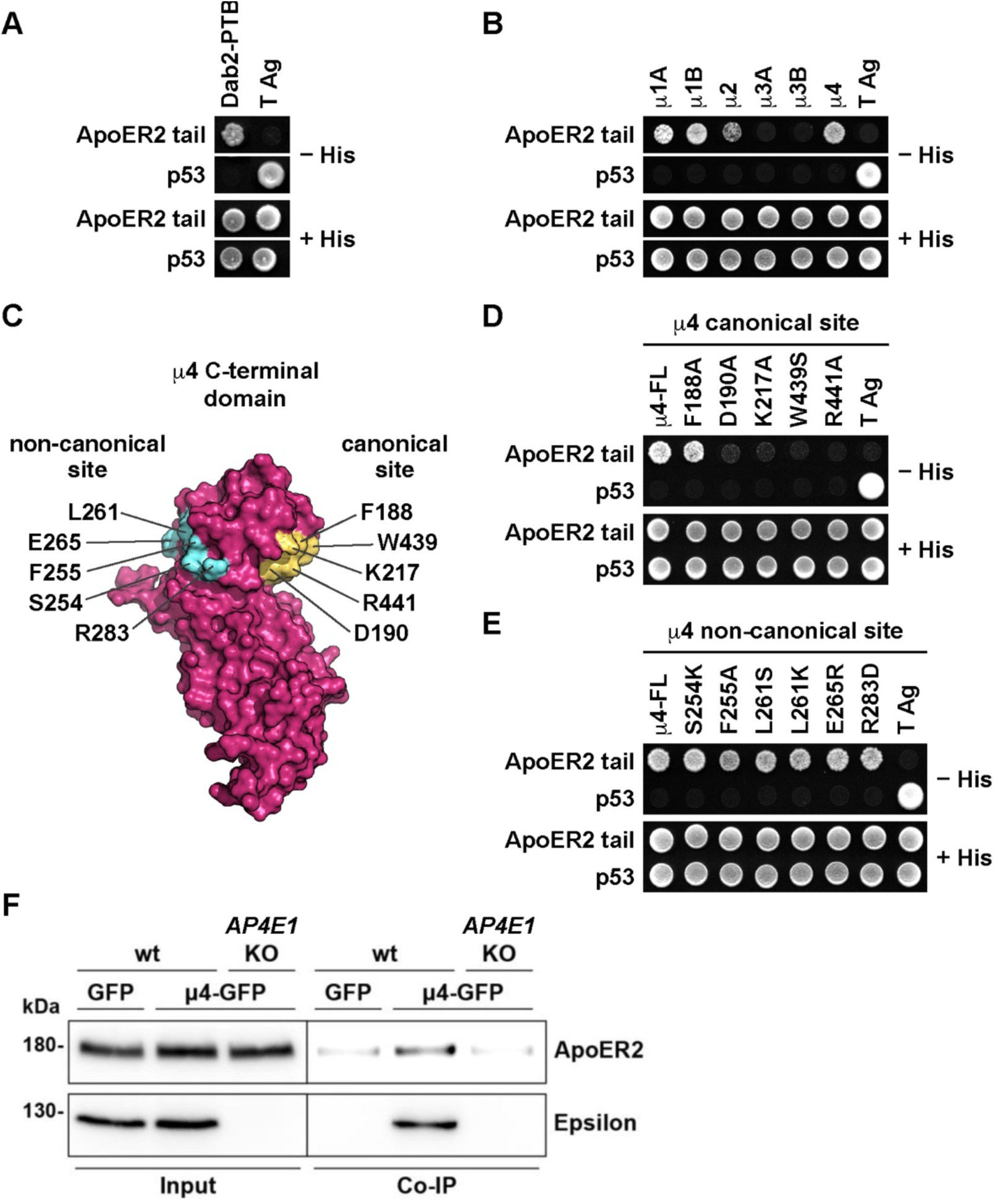
Analysis of the interaction between the ApoER2 cytosolic tail and adaptor proteins. **(A, B)** For Y2H analysis, yeast cells were co-transformed with plasmids encoding Gal4bd fused to wt ApoER2 cytosolic tail (residues 849-963; see Fig. 2A), and Gal4ad fused to Dab2-PTB **(A)** or to the indicated wt µ subunits of the adaptor protein complexes AP-1, AP-2, AP-3, and AP-4 (**B**). (**C**) Surface representation of human μ4 C-terminal domain (pdb entry 3L81) highlighting the position of residues at the canonical and non-canonical sites chosen for the Y2H analysis. The image was prepared with PyMOL Molecular Graphics System, version 2.0.6 Schrödinger, LLC. **(D, E)** Y2H analysis after transformation with plasmids encoding Gal4bd fused to wt ApoER2 cytosolic tail and Gal4ad fused to the indicated mutants of the µ4 subunit of AP-4. **(F)** Wild-type and *AP4E1*-KO HeLa cells were co-transfected with plasmids encoding human HA-tagged ApoER2 and μ4-GFP. In addition, cells were co-transfected with plasmids encoding the receptor with GFP as control. Twenty-four h later, cells were lysed, and cell extracts were immunoprecipitated using GFP-trap. The presence of the receptor was detected by immunoblotting. Notice that the receptor was present in all cell extracts (10 % of input), but was immunoprecipitated only from cells expressing μ4-GFP and the ε subunit of AP-4.

The µ4 subunit of AP-4 has a β-sandwich C-terminal domain that harbors a binding site for a YX[FYL][FL]E motif present in the cytosolic tail of members of the amyloid precursor protein (APP) family (referred to as “non-canonical site”) (Fig. 1 C) (Burgos, Mardones et al. 2010, Ross, Lin et al. 2014). In contrast, the homologous β-sandwich C-terminal domains of the µ subunits from AP-1, AP-2 and AP-4 have a conserved binding site for YXXØ motifs on a face opposite that of µ4 (referred to as “canonical site”) (Fig. 1 C) (Owen and Evans 1998, Mardones, Burgos et al. 2013). The C-terminal domain of µ4 does contain conserved residues at a location similar to that of the canonical site in other µ subunits (Fig.1 C), raising the possibility that it could recognize additional sequence motifs via this site. To determine the requirement of residues in the non-canonical and canonical sites of µ4 for binding to the ApoER2 tail, we performed Y2H assays using µ4 constructs with single-amino acid substitutions at those sites. We found that substitutions of conserved residues in the canonical site (i.e., D190, K217, W439, R441) abolished the interaction (Fig. 1 D). In contrast, substitutions of conserved residues in the non-canonical site did not affect the interaction (Fig. 1 E). ApoER2 is thus the first known cargo recognized by the predicted canonical site on µ4.

To assess if the interaction observed by Y2H can also be detected in mammalian cells, we performed co-immunoprecipitation assays using HeLa cells co-expressing HA-tagged human ApoER2 (HA-ApoER2) and GFP-tagged µ4 (µ4-GFP) or GFP (background control). These assays showed greater co-immunoprecipitation of HA-ApoER2 with µ4-GFP than with GFP (Fig. 1 F). Moreover, the co-immunoprecipitation of HA-ApoER2 with µ4-GFP was reduced to background levels upon KO of the ε (*AP4E1*) subunit of AP-4 (Fig. 1 F), a condition in which the remaining subunits of AP-4 are degraded (Davies, Itzhak et al. 2018, De Pace, Skirzewski et al. 2018). This finding indicated that co-immunoprecipitation of HA-ApoER2 with µ4-GFP requires assembly of µ4 into the AP-4 complex and, therefore, its association with membranes.

The cytosolic domain of ApoER2 lacks YXXØ motifs that could interact with the canonical binding site on µ4 (Fig. 2 A). However, interactions of µ4 with some cargos, such as the δ2 glutamate receptor, depend on Phe residues in the cytosolic domains (Yap, Murate et al. 2003). The human ApoER2 tail contains three Phe residues: Phe-861 (F13, considering the first amino-acid residue after the transmembrane domain as position 1), Phe-897 (F49), and Phe-924 (F76) (Fig. 2 A). We used the Y2H system to test the possible contribution of these residues to the interaction with µ4. We found that mutation of either F13 or F76 to Ala did not affect the interaction, whereas mutation of F49 to Ala abolished it (Fig. 2 B). Further alanine-scan mutagenesis of the segment spanning I46 to D50 (i.e., ISSFD) revealed an additional requirement of I46 for the interaction (Fig. 2 C). Therefore, the interaction of ApoER2 with µ4 depends on the sequence ISS<u>F</u>, in which the Ile and Phe residues (underlined) are essential, and both Ser residues are dispensable. Using purified, recombinant components, we confirmed this interaction by isothermal titration calorimetry (ITC). The synthetic peptide YPAAISS<u>F</u>DR (Fig. 2 D), but not the substituted variant YPAA<u>A</u>SS<u>A</u>DR (Fig. 2 E), bound to a single site on the recombinant µ4 C-terminal domain with *K_d_* = 26.2 ± 6.6 µM. Further Y2H analyses showed that the critical F49 in the ISSF motif could be replaced by a tyrosine residue found at the same position in the mouse ortholog of ApoER2 (NM_001369052.1) (Fig. 2 F). This result was consistent with co-immunoprecipitation experiments showing that mouse ApoER2 also binds AP-4 (Fig. 2 G). Our experiments thus demonstrated that ApoER2 binds to AP-4 through an interaction involving an IXXF/Y motif in the cytosolic tail of the receptor and a canonical binding site on the C-terminal domain of µ4.

**Figure 2.**
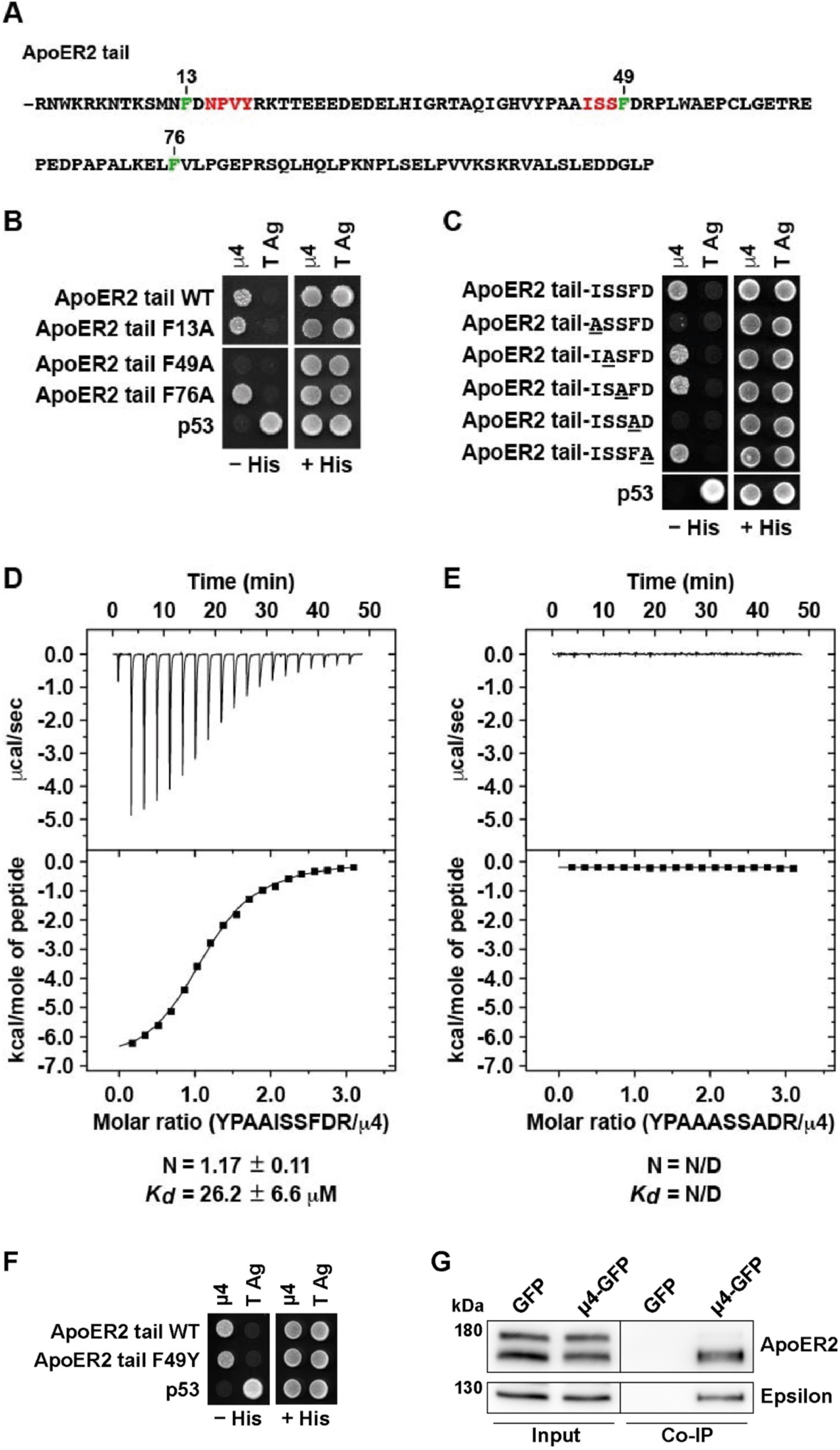
ApoER2 binds via an IXXF/Y motif to a canonical site on the µ4 subunit of AP-4. **(A)** Sequence of the ApoER2 cytosolic tai (residues 849-963), highlighting in green three phenylalanine residues at positions 13, 49 and 76 relatives to R849 as residue 1 of the indicated cytosolic tail sequence. Highlighted in red are an NPXY motif and the sequence ISS adjacent to F49 that is important for interaction to µ4. **(B, C)** Y2H analysis of critical residues on the ApoER2 cytosolic tail for interaction with µ4. Yeast cells were co-transformed with plasmids encoding Gal4bd fused to wt or the indicated mutants of the ApoER2 cytosolic tail, and Gal4ad fused to the wt µ4 subunit of the AP-4 complex. Mouse p53 fused to Gal4bd and SV40 large T antigen (T Ag) fused to Gal4ad were used as controls. Co-transformed cells were spotted onto His-deficient (−His) or His-containing (+His) plates and incubated at 30 °C. **(D-E)** Isothermal titration calorimetry of YPAAISSFDR peptide **(D)** or YPAAASSADR peptide **(E)** with recombinant µ4 C-terminal domain. The *K*_d_ and stoichiometry (N) for the µ4-YPAAISSFDR interaction are expressed as the mean ± SD (n = 3). N/D: not determined. **(F)** Y2H analysis of the effect of substituting tyrosine for F49 in the cytosolic tail of human ApoER2 on interaction with μ4. The analysis was performed as described for panels **B** and **C**. **(G)** Wild-type HeLa cells were co-transfected with plasmids encoding FLAG-tagged mouse ApoER2 and μ4-GFP or GFP as control. Twenty-four h later, cells were lysed, and cell extracts were immunoprecipitated using GFP-trap. The presence of the receptor was detected by immunoblotting. Notice that the receptor was present in both cell extracts (8% of input) but was immunoprecipitated only from cells expressing μ4-GFP.

### 3.2. Recognition of ApoER2 by AP-4 regulates the Golgi and endosomal localization of the receptor and promotes its axonal localization

Next, we sought to determine the importance of the ApoER2–AP-4 interaction for the distribution and trafficking of the receptor. HA-ApoER2-GFP expressed by transient transfection in HeLa cells was distributed among several compartments, including the plasma membrane, cytoplasmic vesicles, and the GC (i.e., an assembly of *cis*, medial and *trans* Golgi cisternae and the TGN) (Fig. 3 A), consistent with trafficking of the receptor through all these compartments. Since AP-4 was previously shown to promote export of cargo proteins from the GC (Burgos, Mardones et al. 2010, Mattera, Park et al. 2017, Ivankovic, Drew et al. 2020, Davies, Alecu et al. 2022), we examined the co-localization of HA-ApoER2-GFP with the GC marker GM130 (Golgi cisternae). Compared to wild-type (wt) HeLa cells, *AP4E1*-KO HeLa cells exhibited increased co-localization of HA-ApoER2-GFP with GM130 as assessed by calculation of Pearson’s correlation coefficients (Fig. 3 A, C). The F49A-mutant ApoER2 that cannot interact with µ4 (Fig. 2 B and C) exhibited a similar co-localization with GM130 in both wt and *AP4E1*-KO cells but increased co-localization relative to the wt receptor (Fig. 3 B, C). Similar results were obtained by examining the co-localization of the wt and F49A mutant ApoER2 with the TGN marker p230 (Fig. S1 A-D). Interestingly, in the absence of AP-4, the endosomal localization of ApoER2 was also modified, showing an increased presence in EEA1-positive endosomes, but not in TfR-and AP-1-positive compartments (Fig. S2 A-G). In contrast to ApoER2-GFP, the transferrin receptor tagged with GFP (TfR-GFP) showed similar Pearson’s correlation coefficients in *AP4E1*-KO and wt cells with respect to Golgi markers and EEA1 (Fig. S3). These experiments thus demonstrated that depletion of AP-4 or mutation of F49A in the ApoER2 cytoplasmic domain increased the steady-state localization of the receptor to the GC, consistent with a role of AP-4 in ApoER2 trafficking from this compartment. Besides, AP-4 seems to negatively regulate the presence of ApoER2 in an EEA1-positive endosomal compartment. A related role for AP-4 in controlling the endosomal localization of cargos was also supported by the increased presence of ATG9A in a TfR-positive compartment in HeLa *AP4E1*-KO (Fig. S4).

**Figure 3.**
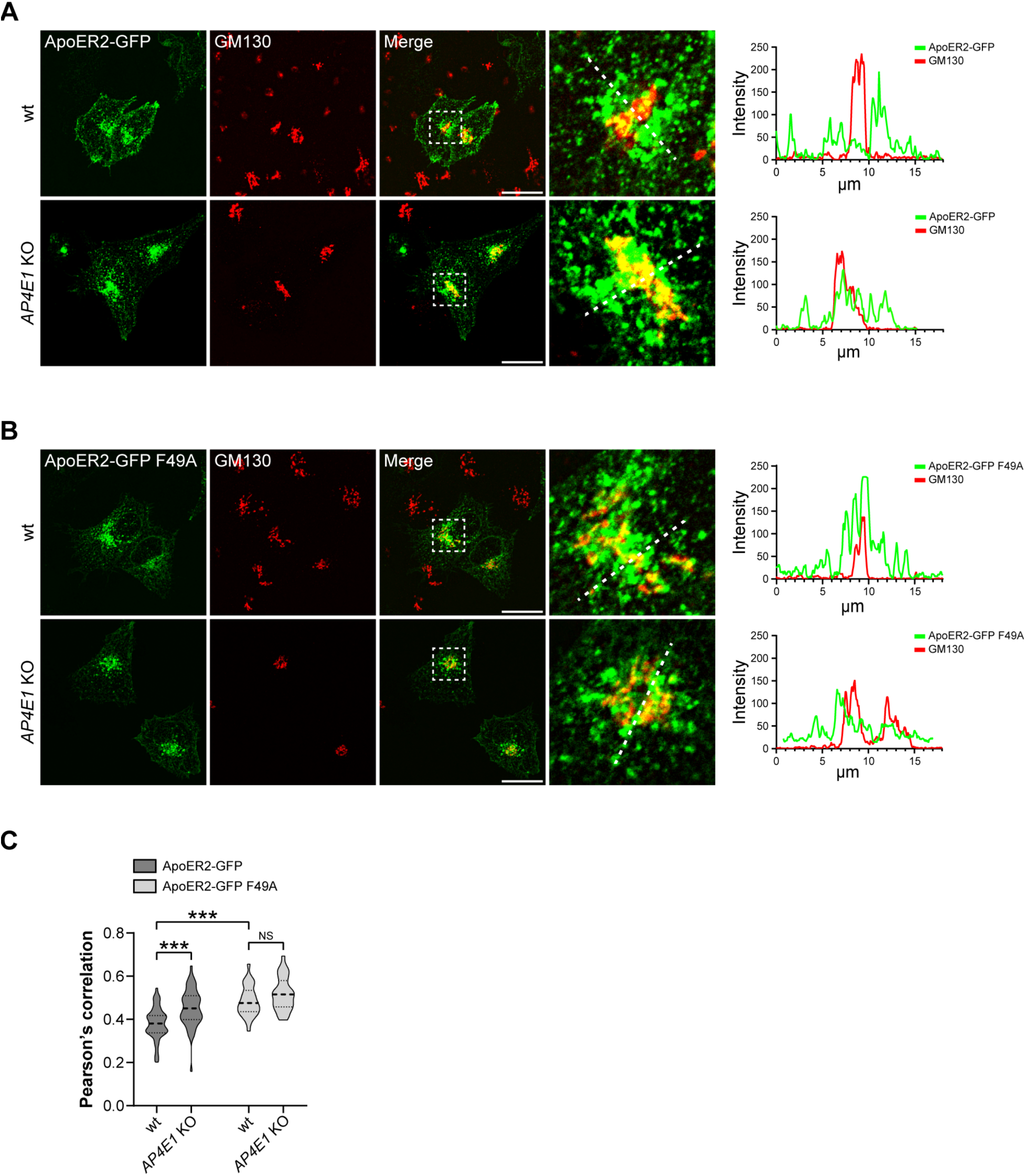
Increased co-localization of ApoER2 with Golgi marker GM130 in *AP4E1*-KO Hela cells. **(A, B)** HeLa cells were transfected with plasmids encoding wt HA-ApoER2-GFP wt (green) **(A)** or HA-ApoER2-GFP F49A (green) **(B)** and immunostained 24 h later with anti-GM130 (red). Scale bar 20 μm. Magnified views of boxed areas are shown on the right. 20 μm-dotted lines were drawn to calculate the intensity profile using Fiji. **(C)** Graphs showing the Pearson’s correlation coefficient obtained from 39-58 cells pooled from 3-4 independent experiments. Statistical significance was calculated by ANOVA with Tukey’s multiple comparisons test, ***p<0.001.

Since ApoER2 is not expressed at detectable levels in HeLa cells, we sought to confirm our results in neurons that endogenously express the receptor and respond to the ligand Reelin (Stockinger, Brandes et al. 2000). We observed that endogenous ApoER2 also exhibited increased co-localization with GM130 in hippocampal neurons from *Ap4e1*-KO relative to wt mice (Fig. 4 A, B). As was found in HeLa cells, the absence of AP-4 did not affect the distribution of ApoER2 in endosomes positive for TfR (Fig. 4C, D). The polarized nature of neurons allowed us to quantify the presence of endogenous ApoER2 in the somatodendritic and axonal domains of *Ap4e1*-KO and wt neurons (Fig. 4E). We observed that the number of ApoER2-containing vesicles within dendrites was similar in *Ap4e1*-KO and wt neurons (Fig. 4 E, F). In contrast, there were significantly fewer ApoER2-containing vesicles in axons from *Ap4e1*-KO relative to wt neurons. These results indicated that the increased presence of ApoER2 in the GC correlated with decreased receptor localization to the axon, but not the dendrites.

**Figure 4.**
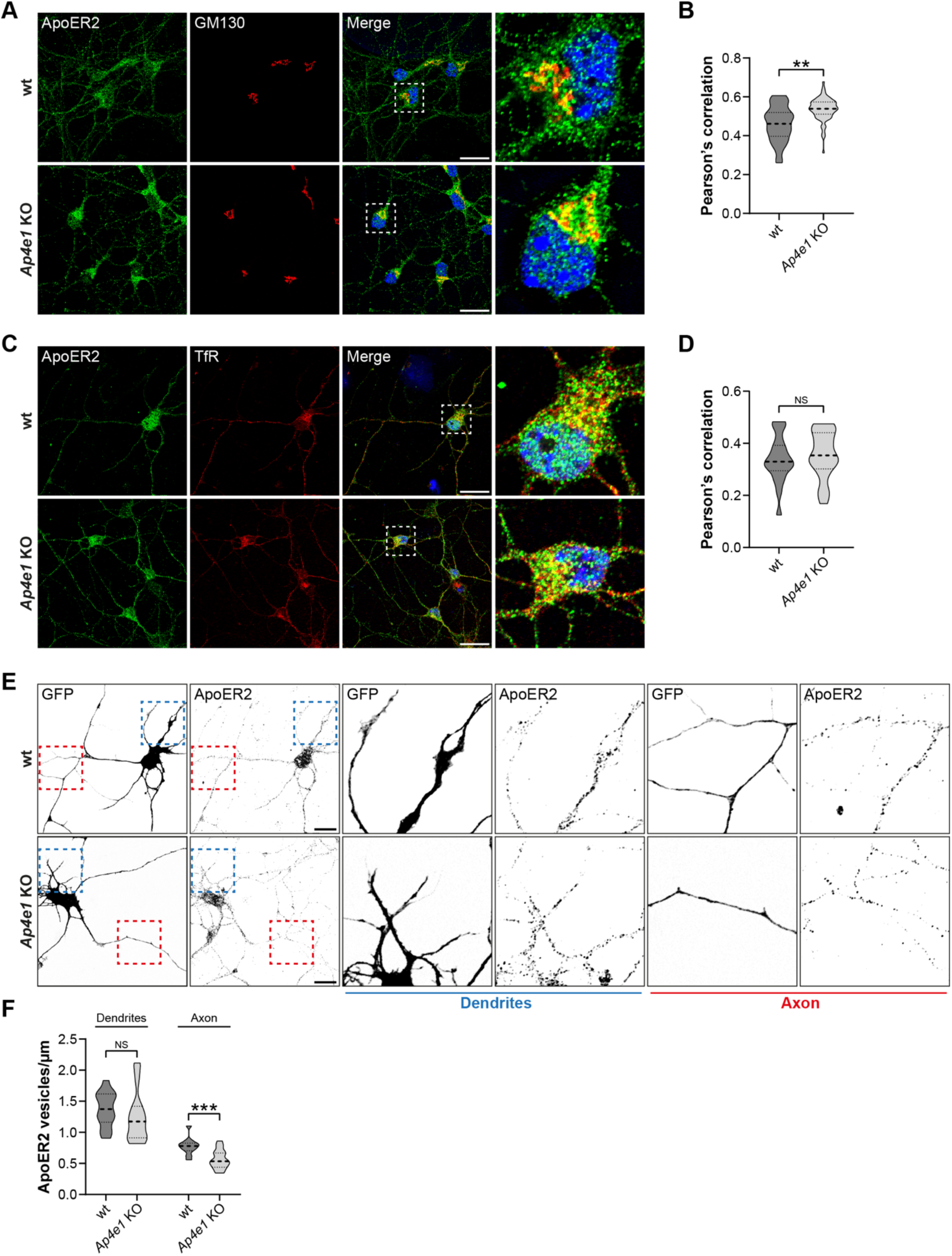
Endogenous ApoER2 shows increased Golgi localization and reduced axonal distribution in *Ap4e1*-KO neurons. **(A)** DIV5 mouse hippocampal neurons from wt and *Ap4e1*-KO mice were immunostained for endogenous ApoER2 (green) and GM130 (red). Magnified views of boxed areas are shown on the right. Scale bar: 20 μm. **(B)** Graph showing Pearson’s correlation coefficient obtained from 36-60 neurons from 3 independent experiments. Statistical significance was calculated using the Mann-Whitney t-test. **p<0.01. **(C)** DIV5 mouse hippocampal neurons from wt and *Ap4e1*-KO mice were immunostained for endogenous ApoER2 (green) and TfR (red). Magnified views of boxed areas are shown on the right. Scale bar: 20 μm. **(D)** Graph showing Pearson’s correlation coefficient obtained from 20 neurons from one experiment. Statistical significance was calculated using the Mann-Whitney t-test. ns: not significant. **(E)** neurons from wt and *Ap4e1*-KO mice were transfected with a plasmid encoding EGFP, fixed 24 h later, and immunostained for endogenous ApoER2. Blue and red boxes show dendrites and axons, respectively. Magnified views of these boxes are in the middle and right panels. Scale bar: 20 μm. **(F)** Number of vesicles per μm was quantified using the analyze particles tool in Fiji. Bar graphs are representative of 12-20 neurons from 2 independent experiments. Statistical significance was calculated using the Mann-Whitney t-test. **p<0.01; ns: not significant.

To complement the above experiments, we evaluated the polarized distribution of transfected HA-ApoER2-GFP (Fig. 5 A, B) or a version without GFP at the C-terminus (HA-ApoER2) (Fig. 5 E, F) in wt and *Ap4e1*-KO hippocampal neurons by calculating the dendrite/axon polarity index (Farias, Cuitino et al. 2012, Guo, Farias et al. 2016). In wt neurons, HA-ApoER2-GFP was more concentrated in the dendrites (labeled for MAP2) than in the axon (Fig. 5 A), as reflected by a polarity index of 2.2 (Fig. 5 B). Similar results were obtained for HA-ApoER2 (Fig. 5 E, F), indicating that the tag’s presence and topological location does not affect the polarized distribution of the receptor. In *Ap4e1*-KO neurons, the wt receptor exhibited reduced presence along the axon, as indicated by the intensity and distribution of the label in *Ap4e1*-KO vs. wt axons (Fig. 5 C, D, and G, H). In dendrites, ApoER2 showed no difference in *Ap4e1*-KO vs. wt neurons (Fig. S5 A-C), similar to what was found with the endogenous receptor (Fig. 4F). In wt neurons, the F49A-mutant exhibited a higher polarity index than the wt receptor and this index did not change upon KO of *Ap4e1* (Fig. 5 B). Accordingly, the axonal intensity and distribution of the mutant receptor was lower than the wt ApoER2 in wt neurons and did not change in *Ap4e1*-KO neurons (Fig. 5C and D). Overall, these results are consistent with the idea that the interaction of ApoER2 with AP-4 preferentially promotes the axonal localization of the receptor.

**Figure 5.**
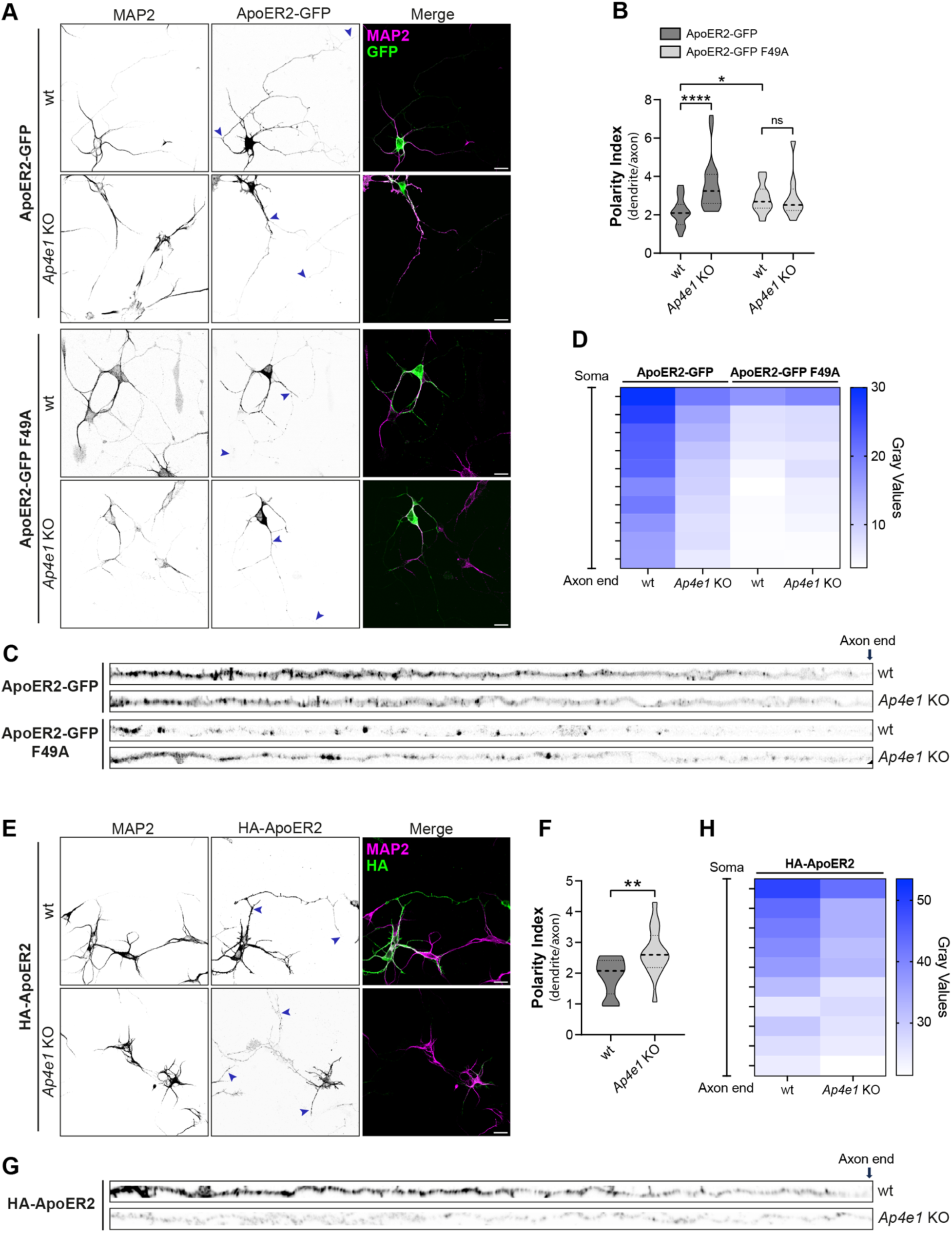
AP-4 promotes axonal localization of ApoER2 in hippocampal neurons. **(A)** wt and *Ap4e1*-KO mouse neurons were transfected at DIV4 with plasmids encoding HA-ApoER2-GFP or HA-ApoER2-GFP F49A (green) and fixed 24 h later. Neurons were immunostained with antibody to MAP2 (magenta) to identify dendrites. Arrowheads show the beginning and end of the axonal segment traced for further analysis. Scale bar: 20 μm. **(B)** The dendrite/axon polarity index of neurons from experiments was calculated as described in Methods from 10-24 neurons in n=4 independent experiments (ApoER2-GFP) or n=2 independent experiments (F49A). Statistical significance was calculated by Mann-Whitney t-test. *p<0.001, **** p<0.0001., ns, not significant **(C)** Images depicting the full length of the axon straightened from the axonal segments in panel A. **(D)** Axonal gray values of GFP signal throughout full-length axons were binned, averaged, and plotted as a heat map to show the spatial distribution of ApoER2 relative to the neuronal soma (see Methods). **(E)** wt and *Ap4e1*-KO mouse neurons were transfected at DIV4 with plasmids encoding N-terminal tagged HA-ApoER2, fixed 24 h later, and stained with anti-HA (green) and anti-MAP2 (magenta). Arrowheads show the beginning and end of the axonal segment traced for further analysis. Scale bar: 20 μm. **(F)** The dendrite/axon polarity index was calculated as described in Methods from 15-17 neurons obtained in n=3 independent experiments. Statistical significance was calculated by Mann-Whitney t-test. *p<0.001. **(G)** Images depicting the full-length of the axon straightened from the axonal segments in panel E. **(H)** Axonal gray values of HA signal throughout full-length axons were binned, averaged, and plotted as a heat map to show the spatial distribution of ApoER2 relative to the neuronal soma (see Methods).

### 3.3. Post-Golgi, AP-4-dependent sorting mechanisms determine the axonal distribution of ApoER2

To directly analyze if the ApoER2-AP-4 interaction mediates ApoER2 export from the TGN, we used a pulse-chase system based on a receptor construct (FM4-HA-ApoER2-GFP) containing four modules of a mutated version of FKBP12 (FM) (Fig. 6 A), which allows for the oligomerization of these proteins in the ER. This aggregation is quickly reversed upon incubation with DD-Solubilizer, a small membrane-permeable molecule (Al-Bassam, Xu et al. 2012) (Fig. 6 B). HeLa cells were transfected with FM4-HA-ApoER2-GFP, and after 12 h of expression, the receptor was released from the ER by adding DD-Solubilizer. FM4-tagged proteins were analyzed by immunoblotting (Fig. 6 C) and fluorescence microscopy (Fig. 6 D) at different times after the addition of DD-Solubilizer. Immunoblot analysis with anti-HA antibody showed a progressive decrease in the molecular weight of the receptor over 30 min, indicative of furin-catalyzed cleavage in the GC (Fig. 6 C). Fluorescent imaging showed an ER reticular pattern of the FM4-HA-ApoER2-GFP before the addition DD-solubilizer (Fig. 6 D), which changed to a perinuclear Golgi distribution, positive for p230 after allowing exit from the ER. After 60 min of release, the receptor co-localization with p230 dropped significantly, because of exit of the receptor from the Golgi (Fig. 6 D and E).

**Figure 6.**
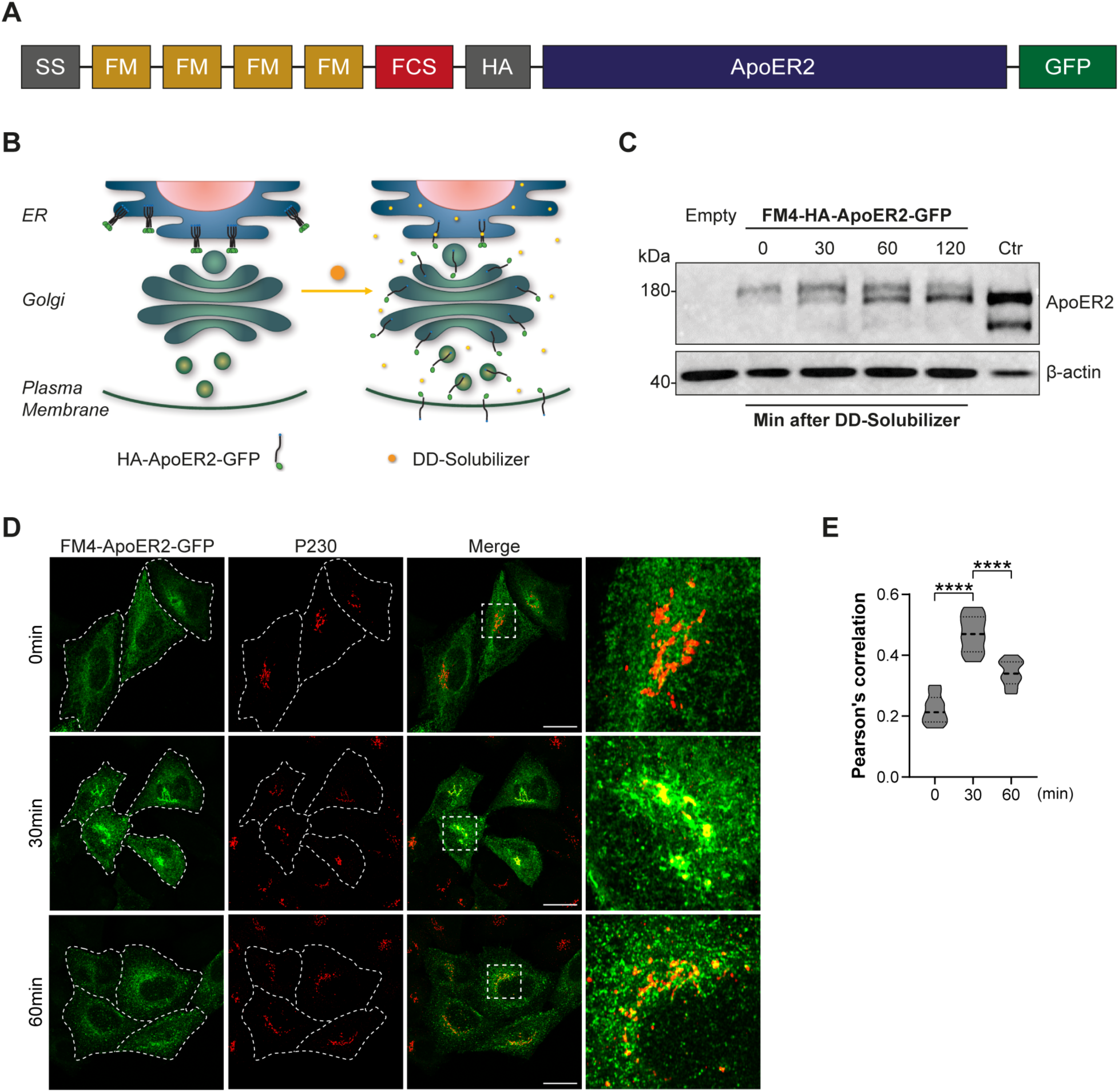
FM4-HA-ApoER2-GFP synchronized release from the ER and its localization to the GC over time in HeLa cells. **(A)** Schematic representation of the construct FM4-HA-ApoER2-GFP. Signal sequence (SS), modified FKBP12 domain (FM), furin cleavage site (FCS), hemagglutinin epitope (HA), apolipoprotein E receptor 2 (ApoER2), green fluorescent protein (GFP) are indicated. **(B)** Schematic representation of the biosynthetic transport of FM4-HA-ApoER2-GFP without (left) and with DD-solubilizer (right). **(C)** Immunoblot detection of the HA epitope (top) and β-actin (bottom) in HeLa cells transfected with plasmids encoding FM4-HA-ApoER2-GFP or the plasmid HA-ApoER2-GFP (ctr lane) without the FM4 module, or untransfected cells (empty lane). Cells extracts were obtained after addition of DD-Solubilizer at the indicated times in min. **(D)** HeLa cells transfected with plasmids encoding FM4-HA-ApoER2-GFP (green) were fixed at the indicated times after addition of DD-Solubilizer. Cells were stained with anti-p230 (red). Dashed white lines delineate the plasma membrane. Scale bar: 20 μm **(E)** Pearson’s correlation coefficient analysis from 35-40 cells pooled from 3 independent experiments for each time point and cell line. Statistical significance was calculated by ANOVA with Tukey’s multiple comparisons test. ****p<0.0001.

To test the possibility that AP-4 regulates the exit of the receptor from the TGN, we compared the exit of wt and F49A mutant FM4-HA-ApoER2-GFP in HeLa cells. Both receptors had the maximum co-localization with Golgi found after 15 min of release (Fig. S6A and B). Surprisingly, there were no significant differences in the presence of newly synthesized receptors (wt vs F49A) in the TGN after 1 h of DD addition. Moreover, when the TGN exit of the wt receptor in the wt vs *AP1E1* KO HeLa cells was compared, there was also no difference (Fig. S6C and D). Altogether, these results indicate that the interaction of ApoER2 with AP-4 does not regulate the exit of the protein from the GC. The increased intensity of ApoER2 in the Golgi area of *AP4E1*-KO cells at steady-state (Fig. 3) could thus reflect an accumulation of the receptor in a neighboring endosomal compartment, which are in close and dynamic contact with the TGN (Fujii, Kurokawa et al. 2020) and cannot be resolved from the TGN by diffraction-limited microscopy.

After analyzing the expression and trafficking of FM4-ApoER2 in HeLa cells, these receptor constructs were used to evaluate if the axonal distribution of the receptor, reduced by the absence of AP-4, depends on an early sorting process, occurring at or immediately at the GC. For this purpose, we performed experiments of synchronized secretory trafficking in DIV6 rat hippocampal neurons transfected with wt and F49A mutant FM4-ApoER2-GFP (Fig. 7). After 16 h of expression, the FM4-tagged receptors were released by the addition of DD-solubilizer and cells were fixed after 30, 60, and 120 min. The polarized distribution of the receptor was determined as previously described (Fig. 5), using the GFP tag to determine the polarity index (Fig. 7 A and B). In the first 30 min, we observed that both constructs readily populated the somatodendritic compartment to similar extents. In contrast, the wt ApoER2-GFP showed enhanced axonal localization at 120 min of release, whereas the F49A mutant was preferentially somatodendritic at the same time of release (Fig. 7A, B). The differential presence of the receptors along the axon was also observed when the intensity and distribution of the label were compared for the wt vs. F49A mutant (Fig. 7C, D). Knowing the general timeline of the polarized trafficking of FM4-ApoER2-GFP in rat neurons, we performed these experiments in mouse wt and *Ap4e1*-KO neurons, fixing the cells after 2 h of release with DD solubilizer (Fig. 8). We observed that the ApoER2 entered the axonal compartment in wt neurons to a greater extent than in the *Ap4e1*-KO neurons (Fig. 8 A-D), consistent with the previous steady-state experiments showing a significant increased polarity index of the receptor in AP-4-deficient neurons (Fig.5). Likewise, the F49A mutant behaved like in rat hippocampal neurons, being preferentially somatodendritic, indicating that AP-4 binding preferentially promotes the axonal localization of ApoER2. Consistently, the localization of the TfR-GFP-FM4 construct remained unchanged in both wt and KO neurons (Fig. S7 A and B). These results suggest that ApoER2 axonal localization takes place after an initial receptoŕs somatodendritic delivery and depends on the interaction of the receptor’s ISSF motif with AP-4. Using this approach, however, we cannot define whether this recognition occurs before or after an eventual arrival to the somatodendritic cell surface.

**Figure 7.**
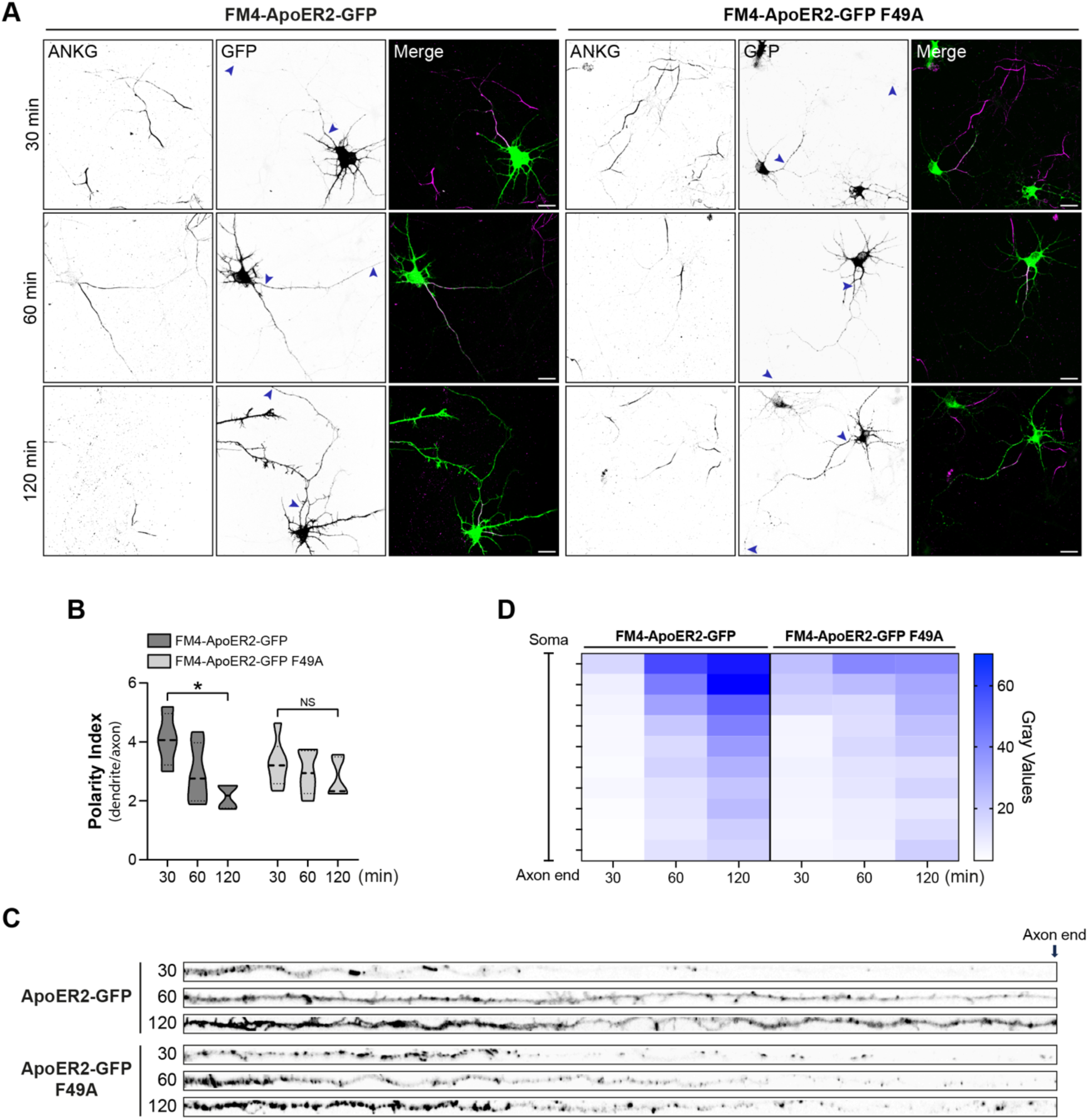
Axonal polarity of ApoER2 is established during post-Golgi biosynthetic trafficking and depends on the receptoŕs F49 cytosolic amino acid residue. **(A)** DIV7 rat hippocampal neurons were transfected with plasmids encoding wt FM4-HA-ApoER2-GFP or FM4-HA-ApoER2-GFP F49A and fixed at the indicated times after addition of DD-Solubilizer. Neurons were stained with anti-ANKG to identify the axon. Arrowheads point to the beginning and end of the axon. Scale bar: 20 μm. **(B)** The dendrite/axon polarity index was calculated as described in Μethods from 7 neurons for each time point. Statistical significance was calculated by Mann-Whitney T-test. *p<0.05. **(C)** Images depicting the full length of the axon straightened from the axonal segments in panel A. **(D)** Axonal gray values of GFP signal throughout full-length axons were binned, averaged, and plotted as a heat map to show the spatial distribution of ApoER2 relative to the neuronal soma (see Methods).

**Figure 8.**
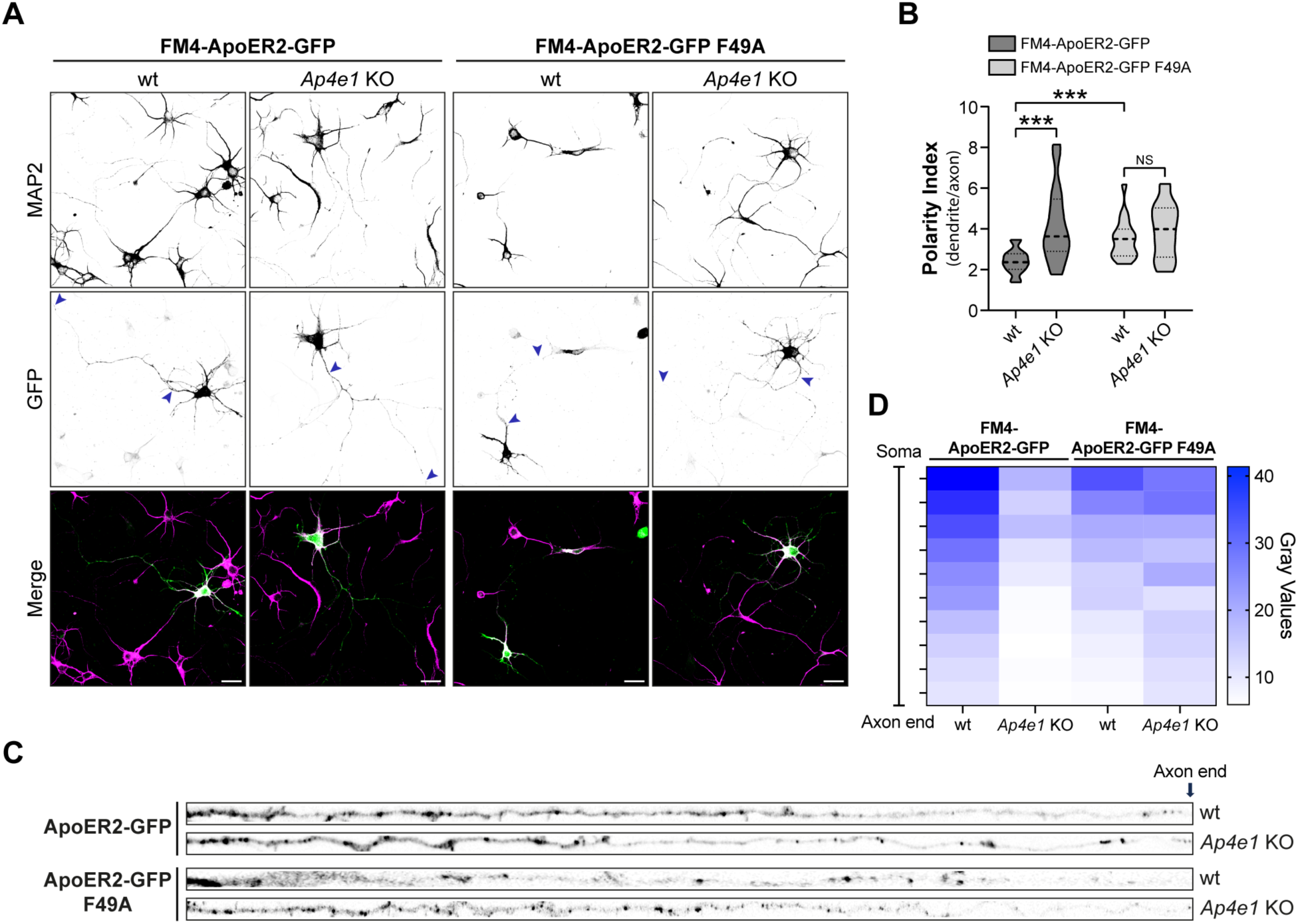
Axonal polarity of ApoER2 is impaired during post-Golgi biosynthetic trafficking in *Ap4e1*-Neurons. **(A)** Wild-type or F49A FM4-HA-ApoER2-GFP were expressed by transfection in DIV4 wt or *Ap4e1*-KO mouse neurons. After 16 h, neurons were treated with DD-Solubilizer for 120 min. Neurons were stained for MAP2 to identify dendrites. Arrows show the beginning and end of the axonal segment traced for further analysis Scale bar: 20 μm. **(B)** The dendrite/axon polarity index was calculated as described in Methods from 17-22 neurons from 3 independent experiments. Statistical significance was calculated using ANOVA with Tukey’s multiple comparisons test *p<0.05. **(C)** Images depicting the full length of the axon straightened from the axonal segments in panel A. **(D)** Axonal gray values of GFP signal throughout full-length axons were binned, averaged, and plotted as a heat map to show the spatial distribution of ApoER2 relative to the neuronal soma (see Methods.)

### 3.4. AP-4 deficiency decreases ApoER2 expression and affects pathways regulated by Reelin signaling

The regulation of ApoER2 trafficking, including its recycling to the plasma membrane, directly impacts the receptor’s function, as evidenced in neurons by the response to one of its ligands, Reelin (Sotelo, Farfan et al. 2014). Therefore, we wondered if the alterations of ApoER2 biosynthetic trafficking due to the disruption of the receptoŕs interaction with AP-4 would affect the receptor total levels, expression at the plasma membrane, and Reelin signaling. These questions were addressed using both hippocampal neurons from wt and *Ap4e1-*KO mice, and wt and *AP4M1*-KO human iPSC-derived cortical-like neurons (i3Neurons) (Fernandopulle, Prestil et al. 2018, Gowrishankar, Lyons et al. 2021).

We observed that the total receptor levels were significantly reduced in DIV14 *Ap4e1-* KO relative to wt mouse hippocampal neurons (Fig. 9 A, C). Likewise, surface biotinylation revealed reduced receptor levels at the plasma membrane of *Ap4e1-*KO relative to wt mouse hippocampal neurons (Fig. 9 B, C). Next, we characterized ApoER2 in the human iPSCs and i3Neurons at various differentiation times up to 30 days. In the wt iPSCs, ApoER2 migrated as several bands, likely resulting from alternative transcript splicing (Fig. 9 D). *AP4M1* KO did not cause detectable changes in either the banding pattern or levels of the receptor (Fig. 9 D). Upon induction of neuronal differentiation, there were time-dependent changes in the banding pattern, although both this pattern and the total levels of the receptor remained equal between wt and *AP4M1*-KO i3Neurons for up to 21 days (Fig. 9 D). At 30 days, however, there was an evident decrease in ApoER2 levels in *AP4M1*-KO relative to wt i3Neurons (Fig. 9 D), like what was found in DIV14 mouse hippocampal neurons (Fig. 9 A). These cultures were repeated several times, and the results consistently showed a significant decrease in total ApoER2 at 30 days (Fig. 9 E, F). The decrease in ApoER2 levels contrasted with the increase in ATG9A levels in *AP4M1*-KO relative to wt i3Neurons (Fig. 9 E, F), as previously shown for this protein in other AP-4-KO cells (Mattera, Park et al. 2017, Davies, Itzhak et al. 2018, De Pace, Skirzewski et al. 2018), and indicating that the drop of ApoER2 is not a consequence of neuronal death. Surface biotinylation showed reduced levels of plasma membrane ApoER2 in *AP4M1*-KO relative to wt i3Neurons at 30 days (Fig. 9 G), as in the mouse neurons (Fig. 9C). ApoER2 mRNA levels did not change in wt vs. KO i3Neurons at 30 days of differentiation, indicating that differences in protein levels were post-transcriptional (Fig. 9 H). To determine when the receptor starts to decrease and if this decrease also takes place preferentially in the axons, as was found in mouse neurons (Fig. 4), we analyzed ApoER2 presence in dendrites and axons at different days of differentiation. After 7 days of differentiation, wt and KO neurons had similar receptor levels (Fig. S8 A and B), but after 14 days of differentiation there was a significant decrease in the intensity of ApoER2 in the axons of KO neurons (Fig. S8C and D), even when the noticeable drop in the total levels, observed by immunoblotting, was detected after 1 month of differentiation (Fig. 9 D). At 30 days of differentiation, we detected a significant decrease in the number of vesicles positive for ApoER2 in axons but not in the soma or dendrites (Fig. 9 I and J). Overall, these results showed that depletion of AP-4 causes an initial reduction in ApoER2 in axons, followed by a drop in more mature neurons and consequently a reduction in the total as well surface ApoER2 available for ligand binding.

**Figure 9.**
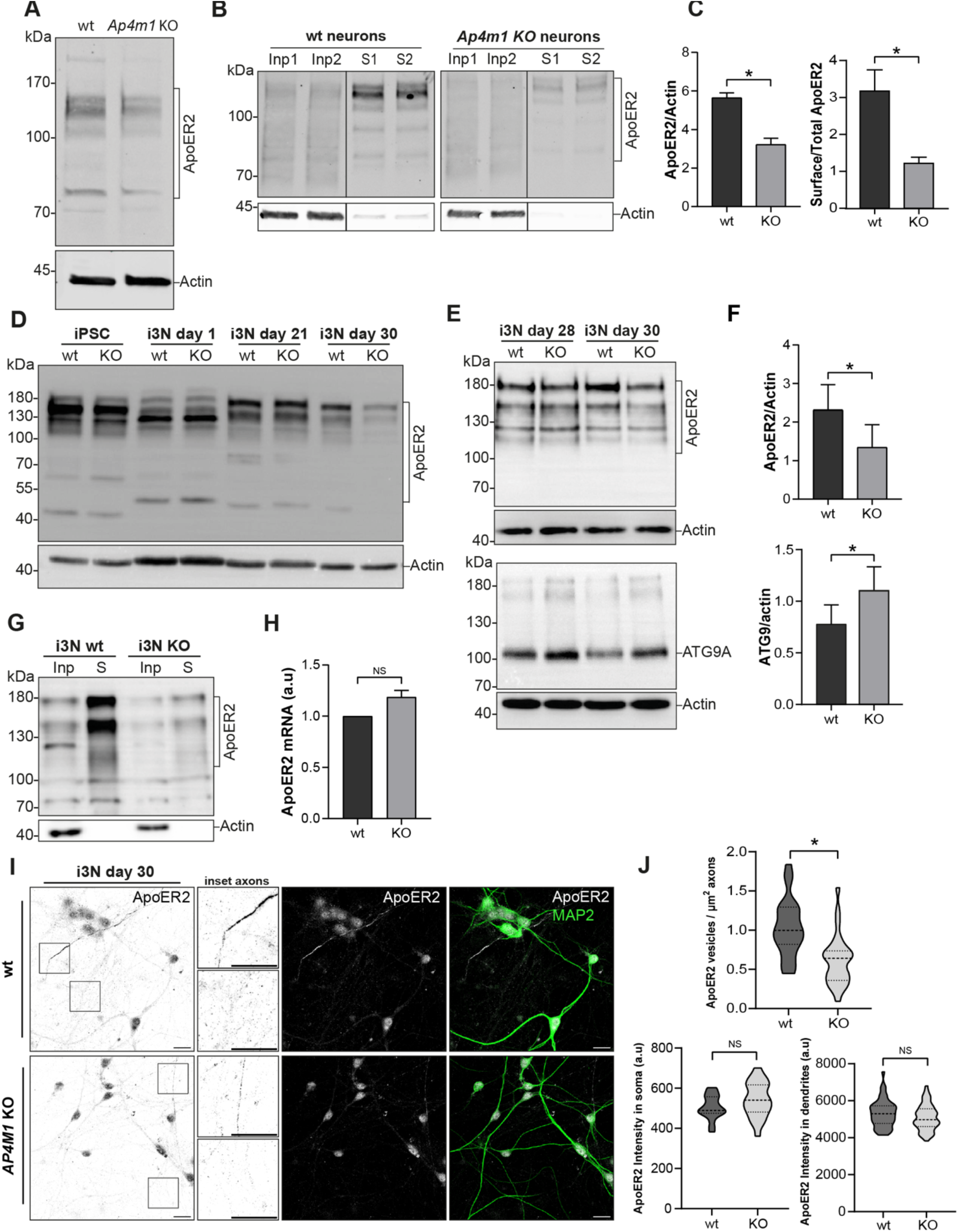
ApoER2 total, axonal and surface levels are reduced in neurons lacking AP-4. **(A)** Immunoblot analysis of DIV12 hippocampal neurons from wt and *Ap4e1*-KO mice showed reduced levels of total ApoER2. **(B)** Cell surface proteins were biotinylated and analyzed by immunoblotting. Input, Inp (5% of the cell extract) and cell surface (S) levels of ApoER2 are shown for wt neurons (left blot) and *Ap4e1*-KO neurons (right blot) **(C)** ApoER2 total levels (n=3) and cell surface with respect to total levels corrected by actin (lower graph) (n=2). Statistical significance was calculated using the Mann-Whitney t-test. *p<0.05. **(D)** Cells extracts from wt and *AP4M1*-KO iPSCs and i3Neurons (i3N) at 1, 21 and 30 days of differentiation were analyzed by immunoblotting for ApoER2 and actin (control). ApoER2 exhibited a differential band pattern through differentiation and a reduction in total levels of the receptor at 30 days in *AP4M1-*KO compared with wt i3Neurons. **(E)** Wild-type and *AP4M1*-KO i3Neurons, differentiated for 28 and 30 days were lysed and analyzed by immunoblotting. Notice that ApoER2 levels were reduced (upper panel) and ATG9A levels increased in *AP4M1*-KO i3Neurons (lower panel). (**F**) ApoER2 and ATG9A total levels (n=5) from i3 Neurons, differentiated for 30 days. Statistical significance was calculated using t-test. *p<0.05. **(G)** i3Neurons differentiated for 30 days were surface biotinylated and levels of surface ApoER2 analyzed by immunoblotting. Notice that *AP4M1*-KO i3Neurons had reduced levels of ApoER2 at the cell surface compared with wt i3Neurons. In all the immunoblots, the positions of molecular mass markers (in kDa) are indicated on the left. **(H)** ApoER2 mRNA levels from wt and *AP4M1*-KO i3Neurons differentiated for 30 days (n=3). Expression levels are normalized to GAPDH expression using the delta-delta Cq method (2-ΔΔCq). Statistical significance was calculated using the Mann-Whitney t-test, ns, not significant. **(I)** Wild-type and *AP4M1*-KO i3Neurons, differentiated for 30 days were stained with anti-ApoER2 (grey) and anti-MAP2 (dendrites, green). Boxes show axons. Magnified images of these boxes are shown next to the panels. Scale bar: 20 μm. **(J)** The number of ApoER2 vesicles per μm^2^ of axons (negative for MAP2) was quantified using the analyze particles tool in Fiji. Upper graph represents 28 axon regions for each phenotype from 2 independent replicates. The lower left graph shows mean fluorescence intensity of ApoER2 in the somatic region, 20 neurons for wt and 25 neurons for *AP4M1-*KO. The lower right graph shows intensity of ApoER2 fluorescence measured in several regions of MAP2-positive dendrites. 50-60 regions of interest analyzed per phenotype. (n=2). Statistical significance was calculated using unpaired t-test. *p<0.05, ns, not significant.

Given the reduction in ApoER2 total levels in neurons lacking the AP-4 complex, we tested if Reelin signaling, and the corresponding neuronal responses, could be affected. To address this question, first, we assessed Reelin-induced dendritic outgrowth, using Sholl analysis of DIV5 hippocampal neurons from wt and *Ap4e1-*KO mice previously treated with 20 nM Reelin or mock-treated for 48 h. We found that this response was not affected in *Ap4e1*-KO compared to wt neurons (Fig. 10 A, B). Further analyses showed that, as expected, Reelin caused an increase in the number of primary dendrites, but this increase was the same in wt and KO neurons (Fig. S9A). Furthermore, we observed an increase in the number of branching points in *Ap4e1*-KO vs. wt neurons, but Reelin treatment did not change the number of branch points in either condition, suggesting that this effect is a consequence of deletion of AP-4 and independent of Reelin (Fig. S9B). As the Reelin-induced dendritic outgrowth depends on AKT activation and mTOR (Jossin and Goffinet 2007), we then compared AKT phosphorylation in wt vs. KO neurons. We found that Reelin addition increased phosphorylated AKT levels to similar extents at 20 and 40 min in wt and *Ap4e1*-KO neurons (Fig. 10 C, D). Similar results were obtained in i3Neurons (Fig. 10 E, F). Thus, these Reelin-dependent effects are not dependent on the expression of AP-4 complex.

**Figure 10:**
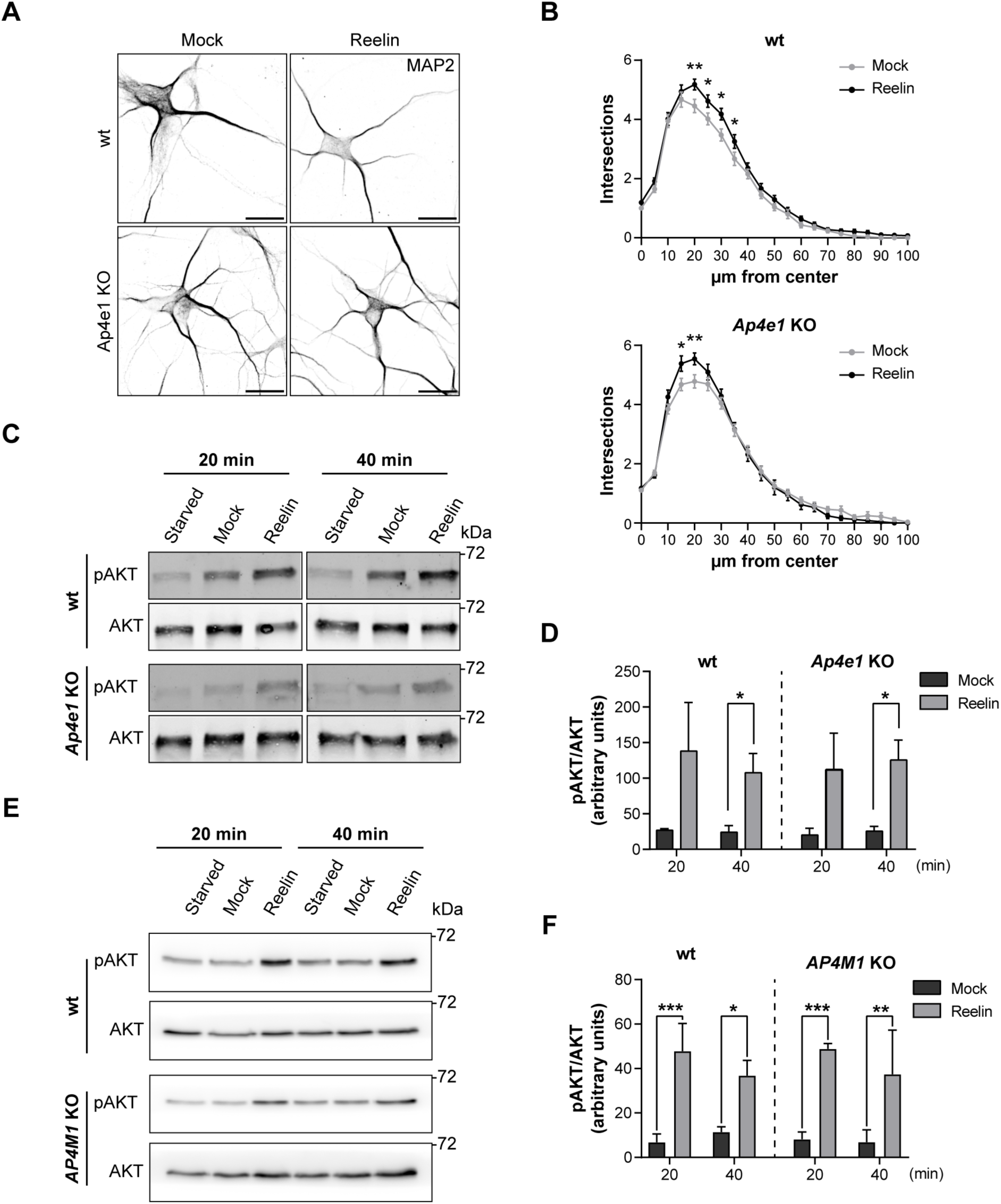
Reelin-induced activation of AKT and dendritic outgrowth are not affected in AP-4-KO neurons. **(A)** DIV5 hippocampal neurons from wt and *Ap4e1-*KO mice were incubated without (mock) or with 10 nM Reelin for 48 h to induce dendritic development. Dendrites were analyzed by immunofluorescence microscopy using an antibody to endogenous MAP2. Scale bars: 20 μm. **(B)** Images from experiments such as that shown in panel A were analyzed using Sholl analysis and the statistical significance was calculated by two-way ANOVA, Sidak’s multiple comparisons test. Results from 50-70 neurons per condition were obtained from 3 independent experiments. *p<0.05, **p<0.01, ***p<0.001. **(C, D)** DIV12-14 wt and *Ap4e1*-KO mouse hippocampal neurons were starved for 4 h and then stimulated with 10 nM Reelin for 20-and 40-min. Neurons were lysed, and the cell extracts subjected to immunoblot analysis to detect pAKT and total AKT. The data from starvation at each time was subtracted from the mock and Reelin conditions to obtain the bar graphs, as specified in Methods. Data are from 3 independent experiments. Statistical significance was calculated with Mann-Whitney t-test-*p<0.05. **(E, F)** Wild-type and *AP4M1-*KO i3Neurons differentiated for 30 days were starved in BrainPhys for 2 h and then stimulated with 10 nM Reelin for 20 and 40 min. Neurons were lysed, and the cell extracts subjected to immunoblot analysis to detect pAKT and total AKT. The data from depletion at each time was subtracted from the mock and Reelin conditions to obtain the bar graphs, as specified in Methods. Data in the graphs correspond to 3 independent experiments. Statistical significance was calculated using Mann-Whitney t-test. *p<0.05, **p<0.01, ***p<0.001. The positions of molecular mass markers (in kDa) in panels C and E are indicated on the right.

Another pathway activated by Reelin involves the phosphorylation of ERK and the downstream activation of CREB (Wang, Xu et al. 2018), as evidenced by CREB translocation to the nucleus (Chen, Durakoglugil et al. 2010). Both responses were measured in i3Neurons after 30 days of differentiation. As expected, Reelin induced the phosphorylation of ERK in the wt neurons, but this response was significantly reduced in the *AP4M1-*KO neurons (Fig. 11 A, B). We also evaluated the activation of CREB following its phosphorylation at serine 133 and its translocation to the nucleus after Reelin treatment for 30 min, as described (Chen, Durakoglugil et al. 2010) (Fig. 11C). The intensity of nuclear fluorescence of pCREB was plotted for each condition (Fig. 11 D). In wt cells, the activation of CREB by Reelin was significant. However, the KO neurons did not show a difference in the nuclear translocation of pCREB in Reelin vs. mock-treated cells. Therefore, the lack of AP-4 reduced the ability of neurons to respond to Reelin in the specific path that includes ERK-pCREB.

**Figure 11:**
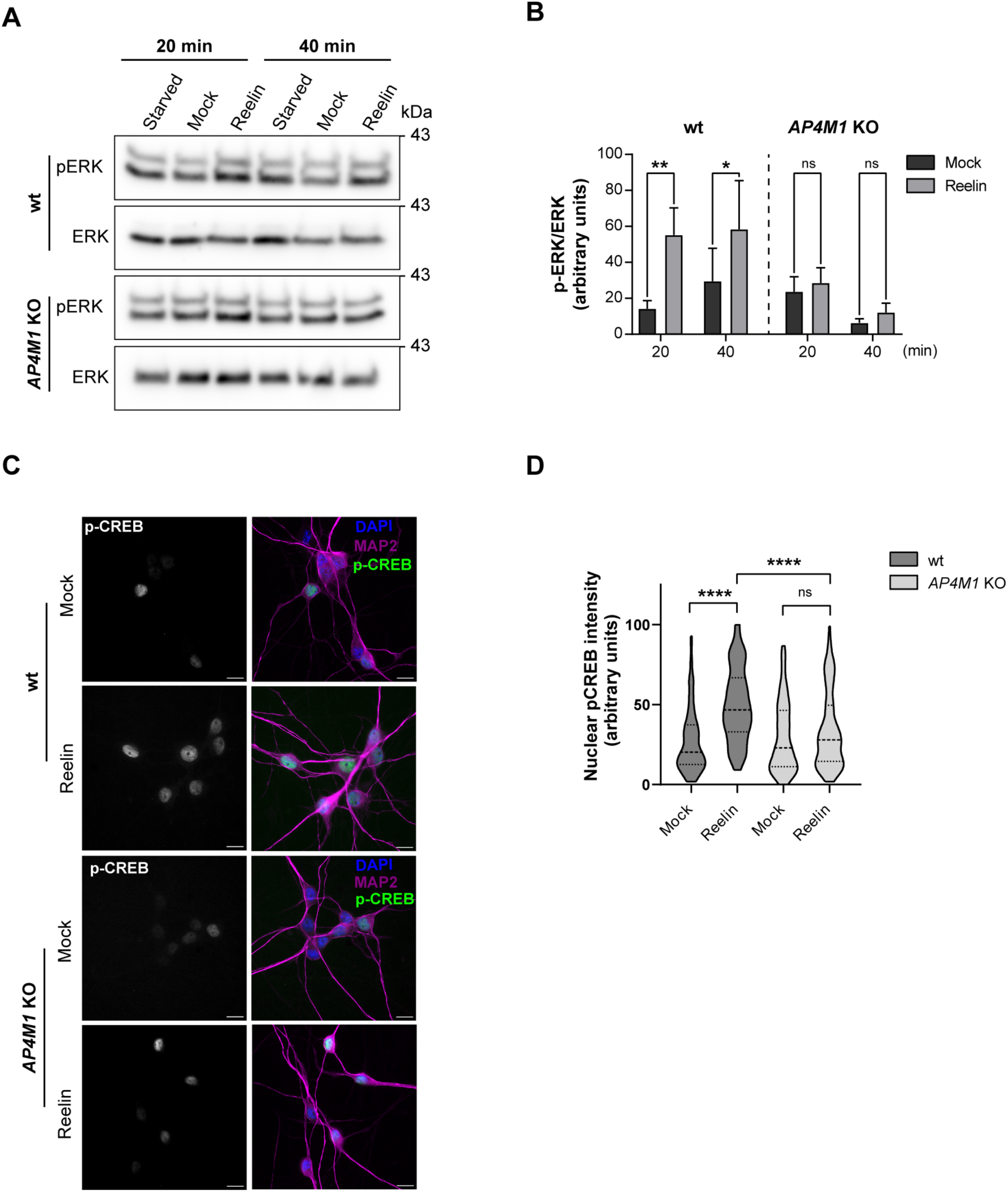
Activation of ERK and CREB induced by Reelin is reduced in neurons lacking AP-4. **(A,B)** Wild-type and *AP4M1-*KO i3Neurons differentiated for 30 days were starved in BrainPhys for 2 h and then incubated without (mock) or with 10 nM Reelin for 20 and 40 min. Neurons were lysed and the cell extracts were subjected to immunoblot analysis to detect pERK and total ERK. The data from depletion at each time was subtracted from the mock and Reelin conditions to obtain the bar graphs, as specified in Methods. Data are from 3 independent experiments. Statistical significance was calculated using Mann-Whitney t-test, * p<0.05, ** p<0.01. **(C)** i3Neurons were starved for 2 h in Hanks’ medium, followed by an additional 30-min incubation in the presence of 10 nM Reelin or mock-conditioned medium. Ser-133–phosphorylated nuclear CREB was detected using a polyclonal phosphorylation-site–specific antibody (red), MAP-2 (green) and nuclei using DAPI (blue). Scale bars: 50 μm. **(D)** Quantification of the nuclear phospho-CREB intensity in neurons treated as in panel C. Data correspond to 2 independent experiments with a total of 223 wt neurons (108 mock, 115 Reelin) and a total of 214 total *AP4M1-*KO neurons analyzed (109 mock, 105 Reelin). Statistical significance was calculated by Kruskal-Wallis’s test with Dunn’s multiple comparisons test, ****p<0.0001.

Finally, we examined a critical aspect of Reelin signaling related to neuronal polarization: the deployment of the Golgi apparatus into the primary dendrite (Meseke, Rosenberger et al. 2013, Caracci, Fuentealba et al. 2019). The experiments were performed in DIV5 mouse hippocampal neurons treated with Reelin. Staining for GM130 (Matsuki, Matthews et al. 2010, Li, Li et al. 2021) showed Reelin-induced Golgi deployment in some of the wt neurons but not in *Ap4e1*-KO neurons (Fig. 12 A and B). Similar results were obtained in i3Neurons (Figure 12 C and D). From these experiments, we concluded that relevant physiological responses of neurons upon Reelin stimulation are compromised in cells devoid of the AP-4 complex.

**Figure 12.**
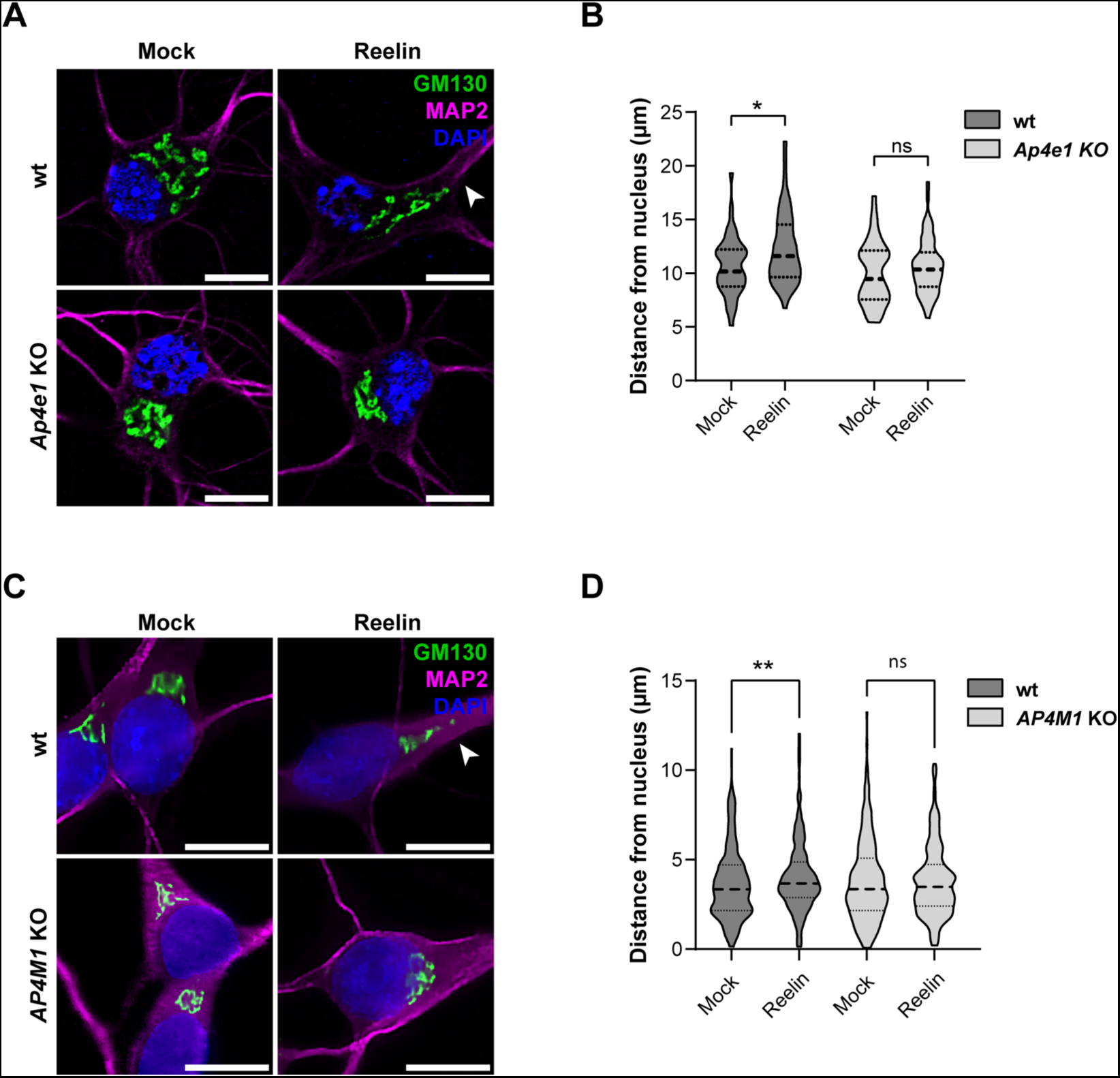
Reelin-induced Golgi deployment is reduced in neurons lacking AP-4. **(A)** Wild-type and *Ap4e*1-KO DIV5 mouse hippocampal neurons treated without (mock) or with 10 nM Reelin for 2 h were stained for GM130 (green), MAP2 (magenta) and nuclei (DAPI, blue). Scale bars: 10 μm. **(B)** Distance from the nucleus to the outermost Golgi was measured using the straight-line tool in Fiji. Statistical analyses were done from 40-60 neurons in 2 independent experiments. Statistical significance was calculated by ANOVA with Tukey’s multiple comparisons test. *p<0.05. **(C)** Wild-type and *AP4M1-*KO i3Neurons were differentiated for 12 days, starved in BrainPhys for 2 h, and treated without (mock) or with 10 nM Reelin for 30 min. Fixed neurons were stained for GM130 (green), MAP2 (magenta) and nuclei (DAPI, blue). Scale bars: 10 μm. **(D)** Distance between the outermost Golgi and the nucleus was measured using a straight-line tool in Fiji. At least 70 cells were measured from 3 independent experiments. Statistical significance was calculated by ANOVA with Tukey’s multiple comparisons test. **p<0.01.

## 4. DISCUSSION

In this work, we determined that ApoER2 binds to AP-4 through an interaction involving an IXXF/Y motif in the cytosolic tail of the receptor and a canonical binding site on the C-terminal domain of µ4. Furthermore, we found that this interaction promotes axonal localization of the receptor as well as neuronal responses to Reelin such as translocation of pCREB into the nucleus and deployment of the GC to the main dendrite. The reduced response to Reelin could be explained by decreased receptor levels, implying that the Reelin signaling pathway, relevant for neurodevelopment and adult brain functions, may be impaired in HSP caused by mutations in subunits of the AP-4 complex.

### 4.1. ApoER2, a new AP-4 cargo that binds to a canonical site on μ4

Our experiments showed that the cytoplasmic domain of the human ApoER2 contains an ISSF motif that is required for binding to μ4, even though it does not fit the previously described YX[FYL][FL]E or YXXØ consensus motifs for μ4 binding (Burgos, Mardones et al. 2010, Bonifacino 2014). Furthermore, the experiments showed that the ApoER2 tail binds to a canonical, μ1/2/3-like binding pocket on the C-terminal domain of μ4, and that a peptide containing this motif binds to the recombinant C-terminal domain of μ4 with a K_d_ = 26.2 ± 6.6 µM, similar to that of the APP tail binding to the non-canonical binding site on μ4 (K_d_ = 29.6 ± 2.4 µM) (Burgos, Mardones et al. 2010). ApoER2 is thus the first AP-4 cargo shown to bind to the canonical binding site on μ4.

It was previously shown that phenylalanine-based motifs could participate in μ4 binding (Yap, Murate et al. 2003). For instance, mutations in any of two hydrophobic/basic motifs (FR) or di-phenylalanine motif (FTF) in the cytosolic tail of the δ2 glutamate receptor resulted in diminished binding to μ4 (Yap, Murate et al. 2003). Most notably, the FTF motifs are contained within an FGSV sequence where a Val residue is also required for binding to μ4 (Yap, Murate et al. 2003), reminiscent of the importance of I46 in the ISSF motif of the ApoER2 tail. However, in contrast to ApoER2, the δ2 glutamate receptor tail was suggested not to interact with the canonical binding site (Yap, Murate et al. 2003).

The role of other phenylalanine-based sorting motifs has been described in the literature. For instance, an IVF motif has been implicated in AP-2 binding to β-arrestins that regulate G-protein-coupled receptors (Burtey, Schmid et al. 2007). In addition, a FI motif is necessary for basolateral sorting of the endoprotease furin, a protein that cycles between the TGN and the plasma membrane (Simmen, Nobile et al. 1999); the same phenylalanine residue is required for the internalization of furin (Stroh, Schafer et al. 1999). Finally, the somatodendritic distribution of the cell adhesion molecule telencephalin, is abrogated by mutation of F905 to Ala in its C-terminal domain (Mitsui, Saito et al. 2005). Phenylalanine residues can sometimes be substituted with tyrosine residues (Bonifacino 2014), as is the case for the somatodendritic/basolateral sorting signals of LRP1 (Donoso, Cancino et al. 2009). In the rat and mouse ApoER2 tails, the Phe residue in the ISSF sequence is replaced by Tyr (ISSY in the rat and ISNY in the mouse receptors). Y2H and coimmunoprecipitation experiments showed that the mouse motif also interacts with μ4. Based on these findings, we can provisionally define the sequence motif that mediates binding of the ApoER2 tail to μ4 as IXXF/Y. Nevertheless, we cannot rule out that additional residues N-or C-terminal to this motif contribute to the interaction. The elucidation of how exactly the cytosolic tail of ApoER2 interacts with the canonical site on μ4 will ultimately require structural analyses of a peptide–μ4 complex.

The relevance of the critical residues identified within the μ4 canonical binding site is supported by the fact that several missense variants reported in HSP patients are clustered within the C-terminal domain. Additionally, residues associated with both the canonical and non-canonical binding sites show significant conservation among vertebrates, making these regions potential hotspots for pathogenic mutations (Sivley, Dou et al. 2018, Gadbery, Abraham et al. 2020). It will be interesting to test if other AP-4 cargo proteins are recognized by this canonical site of the μ4 subunit, including ATG9A (Mattera, Park et al. 2017) and Sortilin 1, a receptor involved in the transport of lysosomal proteins from the TGN to the endolysosomal system that accumulates at the TGN in AP-4 KO neurons (Majumder, Edmison et al. 2022).

### 4.2. Consequences of the lack of interaction between ApoER2 and AP-4 on receptor trafficking

The studies performed in HeLa cells unveiled a role of AP-4 in the Golgi and endosomes. A role of AP-4 in endosomes has been recently suggested in the regulation of AMPA receptor trafficking mediated by tetraspanin TSPAN5 in neurons (Moretto, Miozzo et al. 2023). We found an increase presence of ApoER2 early, but not recycling endosomes in AP-4 deficient cells. In the same way, another AP-4 cargo such as ATG9A was also found increased not only in the Golgi but also in recycling endosomes. The difference regarding the type of endosome in which ATG9A and ApoER2 accumulate in the absence of AP-4 could rely on the particular trafficking characteristics of them; ATG9A has a relevant presence in recycling endosomes (Imai, Hao et al. 2016, Soreng, Munson et al. 2018) but ApoER2 preferentially resides in early endosomes (Sotelo, Farfan et al. 2014). In contrast, TfR which is not an AP-4 cargo, did not change its distribution in the absence of AP-4.

Related to the role of AP-4 in polarized distribution of cargos in neurons, on the one hand our experiments of ApoER2 localization revealed an increase in the dendrite/axon polarity index in *Ap4e1*-KO relative to wt neurons, suggesting a role for AP-4 in the establishment of axonal protein localization. Importantly, these results were not only based on studies of transfected cargo but also relied on experiments with endogenous receptors in both mouse and human neurons, something relevant when the role of adaptor proteins with low level of expression as AP-4 are studied.

AP-4 had already been shown to mediate the polarized distribution of TARPs and by extension AMPA-R in mouse neurons (Matsuda, Miura et al. 2008). However, whereas in our study AP-4 promoted axonal localization, in the case of the TARPs and AMPA-R AP-4 mediated somatodendritic localization (Matsuda, Miura et al. 2008). Like ApoER2, the enzyme DAGLB was also transported into the axon in an AP-4-dependent manner (Davies, Alecu et al. 2022).

In addition, the synchronized trafficking of ApoER2 indicated that its sorting process takes place in the late (from Golgi to plasma membrane) biosynthetic pathway. Hence, the population of FM4-ApoER2 in dendrites 30 min after addition of DD-solubilizer, in both wt and AP-4-KO cells, showed an efficient delivery of the receptor in the biosynthetic pathway; however, in contrast to the wt neurons in which the axonal receptor becomes visible after 30 min, in AP-4 KO neurons little signal was observed in the axon at this time, underscoring the importance of AP-4 in polarized trafficking. Additionally, and consistent with what we observed in AP-4-KO neurons, the ISSF motif of ApoER2 was relevant for axonal localization of the receptor, confirming the role of this motif in polarized trafficking.

To our knowledge, this is the first analysis of the biosynthetic trafficking of an AP-4 cargo, and it is important to note that while previously described cargo have an exclusive somatodendritic localization (e.g. TARPs or Glutamate receptor δ2), ApoER2 is also localized in the axonal domain. The ApoER2 normal polarity appears to be established in a post-Golgi secretory manner with a predominant somatodendritic distribution preceding the appearance in the axonal domain. In contrast, the biosynthetic polarized sorting of other well-known somatodendritic proteins is often incomplete a requires auxiliary mechanisms to maintain its distribution (Guo, Farias et al. 2016). For instance, TfR and GluRs secretory trafficking is seemingly unpolarized with vesicles entering both axons and dendrites in the first minutes after Golgi exit (Al-Bassam, Xu et al. 2012). While some vesicles are excluded in the AIS, other cargo vesicles are actively retrieved from the axon. TfR and glutamate receptors (e.g. GluR1, GluR2, NR2A, NR2B) retrieval is mediated by Rab5 and the FHF complex (Guo, Farias et al. 2016), the latter which also interacts directly with AP-4 (Mattera, Williamson et al. 2020). While the impact of AP-4 deficiency in polarized secretory trafficking of ApoER2 is evident, how this sorting is coupled to post-Golgi trafficking remains unclear.

Nevertheless, it is important to consider alternative routes that could allow the exit of membrane receptors from the GC, ensuring its correct somatodendritic distribution. Recently, it was found that Brefeldin A-Ribosylated Substrate (CtBP1-S/BARS) regulates TGN exit and polarized distribution of membrane receptors such as TfR, L1 and ApoER2 (Gastaldi, Martin et al. 2022). Moreover, our own Y2H screening of adaptor complexes uncovered a direct interaction of ApoER2 with the μ1 subunit of AP-1, which has also been associated to polarized somatodendritic distribution of LRP1 (Donoso, Cancino et al. 2009), and TfR (Farias, Cuitino et al. 2012). While the interaction of multiple adaptor complexes with the same protein cargo has also been described for other proteins such as ATG9A (Imai, Hao et al. 2016, Mattera, Park et al. 2017), the differential contributions of the various APs to polarized trafficking remain to be elucidated.

### 4.3. Roles of ApoER2 in the axon in the context of Reelin signaling

ApoER2 has relevant roles in the somatodendritic domain; specifically, its participation in the regulation of learning and memory processes in postsynaptic terminals has been clearly established in several studies (Beffert, Weeber et al. 2005, Beffert, Durudas et al. 2006, Beffert, Nematollah Farsian et al. 2006). Interestingly, ApoER2 interacts with the post-synaptic scaffold protein PSD 95 (Herz and Chen 2006) and with the polarity protein PAR3 (Pasten, Cerda et al. 2015), in a manner dependent on the presence of exon 18 in humans (exon 19 in mice) (Beffert, Weeber et al. 2005), the same exon encoding the ISSF motif involved in AP-4 binding described in this work.

There is limited information on how ApoER2 functions in the axon and pre-synaptic terminals. ApoER2 co-localizes with pre-synaptic markers and is necessary for local increases in Ca^2+^ and spontaneous neurotransmitter release (Bal, Leitz et al. 2013), underscoring the importance of Reelin in axonal function. Additionally, our group demonstrated that ApoER2 traffics in dorsal root ganglia neurons (DRG) axons in association with TC10 and VAMP7-containing vesicles (Jausoro and Marzolo 2021), and that Reelin stimulates axonal growth in DRGs, further establishing a role for ApoER2 in axonal development and growth.

ApoER2 could also play a role in axonal maintenance and homeostasis. For instance, Reelin has been shown to participate in the specification, elongation, and guidance of axons (Santana and Marzolo 2017, Faini, Del Bene et al. 2021). Defective circuit formation can lead to neurodevelopmental disorders, including intellectual disabilities and neuropsychiatric diseases (Stoeckli 2018). For example, mice that are defective in Reelin signaling have impaired axonal targeting and synapse formation from the entorhinal cortex to the hippocampus (Borrell, Pujadas et al. 2007, Stoeckli 2018, Faini, Del Bene et al. 2021). Similarly, knockdown of ApoER2 in rod bipolar cells in the retina results in the development of swollen and abnormally extended axons. Interestingly, this effect could be recapitulated by the knock-in of ApoER2 with a mutated NPxY motif, which is unable to trigger signaling (Trotter, Klein et al. 2011). These findings suggest that endosomal sorting and Dab1 interaction are essential for axonal development in the retina.

### 4.5. Functional relevance of the ApoER2-AP-4 interaction for Reelin signaling and its implications for HSP

Some of the common phenotypes associated with Reelin and AP-4 deficiencies include defects in axonal growth, guidance, and integrity. In both Reelin and AP-4 deficiencies, there is corpus callosum hypoplasia and cerebellar atrophy, a common finding in various types of HSP, including AP-4-deficiency syndrome (Ebrahimi-Fakhari, Cheng et al. 2018), as well in Lissencephaly 2 (caused by mutations in the *RELN* gene encoding Reelin) (Hong, Shugart et al. 2000). Furthermore, axonal degeneration is observed both in vitro and in vivo in neurons with KO of AP-4 subunits, further suggesting that AP-4 is necessary for axonal maintenance (Matsuda, Miura et al. 2008, De Pace, Skirzewski et al. 2018, Ivankovic, Drew et al. 2020). Additionally, GABAergic interneurons are the primary Reelin-secreting cells in post-natal and adult brains (Pesold, Impagnatiello et al. 1998). This neuronal subtype readily populates the corpus callosum, playing an essential role in interhemispheric connection (Pesold, Impagnatiello et al. 1998). This process has been recently associated with Reelin signaling (Hafner, Guy et al. 2021),suggesting that local Reelin secretion could participate in the maintenance of axonal tracks in the corpus callosum.

Besides decreased axonal localization we observed significantly decreased total and surface protein levels of ApoER2 in AP-4-KO neurons, not explained by diminished mRNA levels. These changes are likely to directly influence the downstream activation of components of the Reelin pathway. However, it is unknown if the local activation of Reelin signaling at the axon or at the somatodendritic domain would affect differentially the components of the pathway. To ascertain the status of Reelin signaling in AP-4-KO cells, we evaluated known phenotypic effects of downstream activation targets.

First, we analyzed AKT phosphorylation, as Reelin directly activates the PI3K-AKT-mTOR axis to promote the growth and branching of developing dendrites (Jossin and Goffinet 2007). Although there is an important body of information regarding the role of Reelin in dendritic development, the role of AP-4 in this process is less clear and seems to depend on the developmental stage of cultured neurons (Behne, Teinert et al. 2020, Ivankovic, Drew et al. 2020). Interestingly, we found dendritic development triggered by Reelin was not affected in AP-4-KO neurons even when the axonal receptor was decreased already at DIV4 in the KO mouse hippocampal neurons (Fig.4). We did not determine Reelin-induced dendritic outgrowth in human cortical neurons, but at the time we could have performed the experiment (after 14 days of differentiation), the receptor was also reduced in axons (Fig. S8). On the other hand, the activation of AKT triggered by Reelin was similar in wt and AP-4-KO neurons even though the total receptor levels were evidently reduced at DIV14 in mouse neurons and after 28-30 days of differentiation in human i3Neurons respectively. In this regard, it is interesting to highlight that we found a specific reduction of ApoER2 in axons and therefore, it is possible that the activation of AKT could take place only in the dendrites where the receptor levels are likely not reduced. Additionally, it could be argued that there is a minimal level of ApoER2 required to respond and activate this branch of the pathway, that a splice-variant of the receptor that is not a cargo of AP-4 is the main responsible for this process, or that VLDL-R, the other Reelin receptor that we did not measure, compensates for the reduction in ApoER2. For instance, in ApoER2-KO neurons, VLDL-R is sufficient to induce phosphorylation of AKT and DAB1 upon addition of Reelin, and signaling activity is only abrogated when both receptors are absent (Beffert, Morfini et al. 2002).

Second, we evaluated Reelin-induced ERK and CREB phosphorylation and found that both modifications were significantly diminished in AP-4-KO neurons. CREB is a transcription factor that is activated downstream of ERK and that has been consistently associated to synaptic plasticity, fulfilling an essential role in memory and learning processes (Benito and Barco 2010). It has been previously shown that Reelin treatment enhances CREB phosphorylation and translocation to the nucleus (Chen, Durakoglugil et al. 2010), and that CREB activates gene expression by forming a complex with the intracellular domain of ApoER2 (Telese, Ma et al. 2015). As mentioned above, the roles of Reelin and ApoER2 in synaptic plasticity are well established (Wasser and Herz 2017). Nevertheless, how synapses are affected in AP-4-deficient neurons has not been addressed. For instance, it is known that AP-4 deficiency leads to missorting of AMPA-R to the axon (Matsuda, Miura et al. 2008) and that sustained synaptic activity induced by tetrodotoxin treatment significantly increases *Ap4m1* RNA levels in wt neurons (Steinmetz, Tatavarty et al. 2016). Seizure disorders have also been documented in AP-4-deficiency syndrome patients (Ebrahimi-Fakhari, Behne et al. 2018), which suggests a potential involvement of synaptic connectivity in the onset of this disease.

Third, we looked at Reelin-induced deployment of the GC, finding that this response was almost absent in AP-4-KO neurons. The mechanisms responsible for GC deployment are mostly unknown but include the activity of the STK25 kinase (Matsuki, Matthews et al. 2010). Reelin activates Golgi deployment via the GTPases Rap1 and Cdc42 (Jossin and Cooper 2011, Meseke, Rosenberger et al. 2013), in a process that is required for neuron polarization and migration during development (Santana and Marzolo 2017). This is a cell-autonomous mechanism thought to underlie the biogenesis of discrete Golgi outposts within developing neurites (Caracci, Fuentealba et al. 2019). In this work, we show that Golgi deployment is also conserved in human iPSC-derived neurons, a finding that was recently published while we were preparing this manuscript (Wang, Cho et al. 2023). It remains to be determined if this phenomenon also occurs in human brain and if it has role in human neurodevelopment, as there is increased interest in the role of the GC in disease (Rasika, Passemard et al. 2018, Boecker, Olenick et al. 2020, El Ghouzzi and Boncompain 2022). It is of note that basal activation levels of downstream targets (e.g., pERK, pAKT, pCREB) in untreated cells are mostly unchanged between WT and AP-4-KO neurons and that it is only upon the addition of Reelin that these defects become readily apparent.

Reelin expression had been previously detected by RT-qPCR analysis in human iPSC-derived neurons (Wang, Ward et al. 2017). Nevertheless, here we address for the first time in this model the functional role of ApoER2 as a membrane-bound signaling receptor and its downstream activity triggered by Reelin. Given the growing importance of Reelin signaling in neuronal development and disease (Santana and Marzolo 2017), as *RELN* is considered an ASD high-confidence gene (Scala, Grasso et al. 2022) and a neuroprotector in AD (Lopera, Marino et al. 2023), the use of human iPSC-derived neurons will be instrumental in the development of novel therapies targeting this signaling pathway. Regarding AP-4-associated HSP, the differences in severity of mouse models with patients carrying mutations in AP-4 subunit genes (Matsuda, Miura et al. 2008, De Pace, Skirzewski et al. 2018, Ivankovic, Drew et al. 2020), warrants the use of novel human cellular models which could better explain this differential physiology. Furthermore, protein trafficking studies in i3Neurons also show the potential to shed light on differences among human and mouse neuronal models (Boecker, Olenick et al. 2020, Wang, Daniszewski et al. 2023).

## 5. CONCLUSIONS

Our work reveals that the plasma membrane receptor ApoER2 is a cargo of the HSP-associated AP-4 complex. The direct interaction between these two components impacts the receptor’s expression and polarized distribution, ultimately influencing the primary function of ApoER2 as a neuronal signaling receptor. The diminished response to Reelin in AP-4-deficient neurons could have profound consequences for early neurodevelopment. Neurons require the ability to sense environmental cues to polarize, migrate and/or differentiate, establishing their position along specific brain regions and their synaptic connectivity with neighboring cells. Further studies of the Reelin-ApoER2 signaling pathway will deepen our understanding of the developmental defects observed in AP-4-deficiency syndrome, paving the way towards novel therapeutic options.

## ACKNOWLEDGMENTS

We thank Dr. Alfredo Cáceres (Centro de Investigación en Medicina Traslacional Severo R. Amuchástegui (CIMETSA), Instituto Universitario de Ciencias Biomédicas de Córdoba (IUCBC), Córdoba, Argentina) for providing plasmids encoding TfR-GFP and pFM4-GFP TfR, and Laura Gastaldi (Instituto Universitario de Ciencias Biomédicas de Córdoba (IUCBC), Córdoba, Argentina) for cloning FM4-HA-ApoER2-GFP. We also thank Dr. Enrique Rodriguez-Boulan (Margaret Dyson Vision Research Institute, Weill Cornell Medical College. USA) for the gift of pFM4-GFP type I, Rosalba Escamilla for her technical support in starting the culture of i3Neurons and performing preliminary experiments, David Necunir Ibarra for designing Graphical Abstract and Figure 6B, the Advanced Microscopy Facility UC and the CIBEM animal facility UC, Dr. Rui Jia (NIH) for the gift of *AP4M1*-KO iPSCs. This work was supported by Fondecyt Regular 1200393 to MPM, CONICYT-PCHA/National Graduate Program/2016-21160410 awarded to MC, CONICYT-PCHA/National Graduate Program/2017-21171004 awarded to LMF, Agencia Nacional de Investigación y Desarrollo (ANID) 2019-21190369 to CA and 2021-21211557 to HP, EQM140116 from Fondo de Equipamiento Científico y Tecnológico of Chile (FONDEQUIP; http://www.conicyt.cl/fondequip), and the Intramural Program of the *Eunice Kennedy Shriver* National Institute of Child Health and Human Development, National Institutes of Health (ZIA HD001607 to JSB).

## AUTHOR CONTRIBUTIONS

GAM, MC, RdP and M-PM: Conceived and designed the experiments. MC, PF, GAM, CA-G, HP, LMF, NS, VAC, TPP and MPM: Performed experiments. JSB, MW, M-PM and GAM: provided essential reagents. MC, GAM, CA-G, HP, LMF, PF, RdP, JSB and M-PM: Analyzed the data. M-PM: Supervised the project, secured the funding. M-PM: Writing-Original Draft. M-PM: Writing. M-PM, GAM, MC, RdP and JSB: Review and editing.

## Declaration of generative AI and AI-assisted technologies in the writing process

During the preparation of this work the author(s) used Grammarly to improve readability and language. After using this tool/service, the author(s) reviewed and edited the content as needed and take(s) full responsibility for the content of the publication.

