## Supplementary Tables and Figures for "The Reelin Receptor ApoER2 is a Cargo for the Adaptor Protein Complex AP-4: Implications for Hereditary Spastic Paraplegia"

| Table S1. Antibodies used in this study | | | | |
|---|---|---|---|---|
| *Immunofluorescence* | | | | |
| **Antibody** | **Host** | **Catalog Number** | **Vendor** | **Dilution** |
| ATG9A | Rabbit | ab108338 | Abcam | 1:250 |
| GM130 | Mouse | 610822 | BD Biosciences | 1:500 |
| Ankyrin-G | Goat | sc31778 | Santa Cruz | 1:50 |
| p230 | Mouse | 611280 | BD Biosciences | 1:500 |
| MAP2 | Chicken | ab5392 | Abcam | 1:500 |
| MAP2 | Rabbit | sc-20172 | Santa Cruz | 1:500 |
| MAP2 | Chicken | PA1-16751 | Thermo Fisher | 1:5000 |
| HA | Chicken | ab3254 | Millipore | 1:250 |
| ApoER2 | Rabbit | A3481 | Sigma-Aldrich | 1:500 |
| phospho-CREB Ser133 | Rabbit | 9198 | Cell Signalling | 1:1500 |
| TfR | Rabbit | ab84036 | Abcam | 1:500 |
| EEA1 | Rabbit | 3288S | Cell Signalling | 1:300 |
| γ-adaptin | Mouse | A4200 | Sigma-Aldrich | 1:1000 |
| Alexa Fluor 488 anti-rabbit IgG | Donkey | A21206 | Thermo Fisher | 1:1000 |
| Alexa Fluor 546 anti-rabbit IgG | Donkey | A10040 | Thermo Fisher | 1:1000 |
| Alexa Fluor 488 anti-mouse IgG | Donkey | A21202 | Thermo Fisher | 1:1000 |
| Alexa Fluor 555 anti-mouse IgG | Donkey | A31570 | Thermo Fisher | 1:1000 |
| Alexa Fluor 488 anti-chicken IgG | Goat | A11039 | Thermo Fisher | 1:1000 |
| Alexa Fluor 647 anti-chicken IgG | Goat | A21436 | Thermo Fisher | 1:1000 |
| Alexa Fluor 647 anti-goat IgG | Donkey | A21447 | Thermo Fisher | 1:1000 |
| *Western Blot* | | | | |
| ApoER2 | Rabbit | A3481 | Sigma-Aldrich | 1:3000 |
| Phospho-Akt (Ser473) | Rabbit | 9271S | Cell Signalling | 1:3000 |
| Akt | Rabbit | 9272S | Cell Signalling | 1:3000 |
| Phospho-p44/42 MAPK (Erk1/2) (Thr202/Tyr204) | Rabbit | 9101S | Cell Signalling | 1:1000 |
| HA | Rabbit | C29F4 | Cell Signalling | 1:1000 |
| ERK (pan ERK) | Mouse | 610124 | BD Biosciences | 1:2000 |
| ε-adaptin | Mouse | 612019 | BD Biosciences | 1:1000 |
| ATG9A | Rabbit | ab108338 | Abcam | 1:1000 |
| HRP-Anti-beta Actin | Mouse | ab49900 | Abcam | 1:5000 |
| HRP-conjugated anti-rabbit IgG | Goat | 111-035-144 | Jackson ImmunoResearch | 1:5000 |
| HRP-conjugated anti-mouse IgG | Goat | AP124P | Sigma-Aldrich | 1:5000 |

| Table S2. Primers used in this study | |
|---|---|
| *Cloning of ApoER2-HA into pFM4 GFP* | Fw GACTAAGCTTTACCCGTACGACGTCCCGGAC<br>Rv CTGAGTCGACTGGGGTAGTCCATCATCTTCAAG |
| *FM4-HA-ApoER2-GFP mutagenesis F49A* | Fw AATCAGCAGCGCTGATCGCCCAC<br>Rv GCTGCAGGATAGACATGG |
| *pBridge ApoER2-FL and HA-ApoER2FL mutagenesis* | |
| Ile-894/Ala (46)* | Fw CTATCCTGCAGCAGCCAGCAGCTTTGATCGC<br>Rv GCGATCAAAGCTGCTGGCTGCTGCAGGATAG |
| Ser-895/Ala (47)* | Fw CCTGCAGCAATCGCCAGCTTTGATCGCCCAC<br>Rv GTGGGCGATCAAAGCTGGCGATTGCTGCAGG |
| Ser-89/Ala (48)* | Fw GCAGCAATCAGCGCCTTTGATCGCCCAC<br>Rv GTGGGCGATCAAAGGCGCTGATTGCTGC |

| | |
|---|---|
| Phe-897 /Ala(49)* | Fw GCAATCAGCAGCGCTGATCGCCCACTG<br>Rv CAGTGGGCGATCAGCGCTGCTGATTGC |
| Asp-898/Ala (50)* | Fw GCAATCAGCAGCTTTGCGCGCCCACTGTGGG-<br>Rv CCCACAGTGGGCGCGCAAAGCTGCTGATTGC |
| Ile-894 /Ala Asp-898/Ala | Fw CTATCCTGCAGCAGCCAGCAGCGCTGATCGC<br>Rv GCGATCAGCGCTGCTGGCTGCTGCAGGATAG |
| *Cloning of AP4M1 into pEGFP* | Fw CACACAGAATTCGCCACCATGATTTCCCAATTCTT<br>Rv CACACAACCGGTACACCTGATCCAGAGCCTCC |
| *PCR genotyping for AP4M1* | Fw CGAAATTGGCGCGAAACGTC<br>Rv CCCAAGTGGAATCTACCAATGAAC |
| *qPCR ApoER2* | Fw GTGGCACTAGATGTGGAAGT<br>Rv GTGCAACTGCTCGTCAATG |
| *qPCR GAPDH* | Fw TCATCAGCAATGCCTCCTG<br>Rv GGCCATCCACAGTCTTCTG |
| *Aminoacidic position relative to the first amino acid of ApoER2 cytoplasmic tail. | |

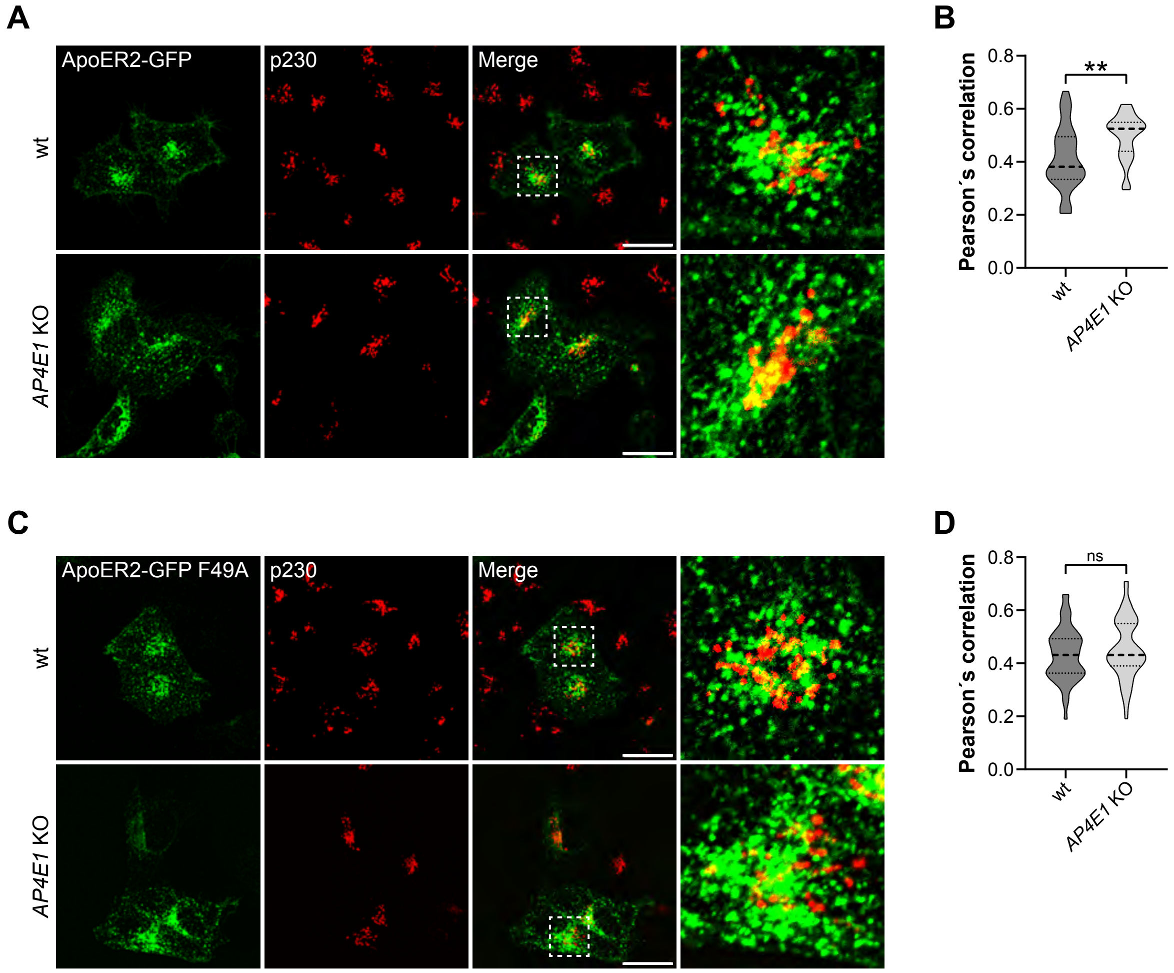

**Figure S1. Increased co-localization of wt ApoER2, but not F49A mutant, with TGN marker p230 in *AP4E1*-KO HeLa cells. (A, C)** HeLa cells were transfected with plasmids encoding wt ApoER2-GFP (green) **(A)** or **(C)** ApoER2-GFP F49A (green) and immunostained 24 h later with p230 (red). Scale bar 20 µm. Magnified views of boxed areas are shown on the right. **(B, D)** Graphs showing the Pearson's correlation coefficient obtained from 36-57 cells pooled from n=3-4 experiments. Statistical significance was calculated by Mann Whitney t-test. **p<0.01, ns not significant.

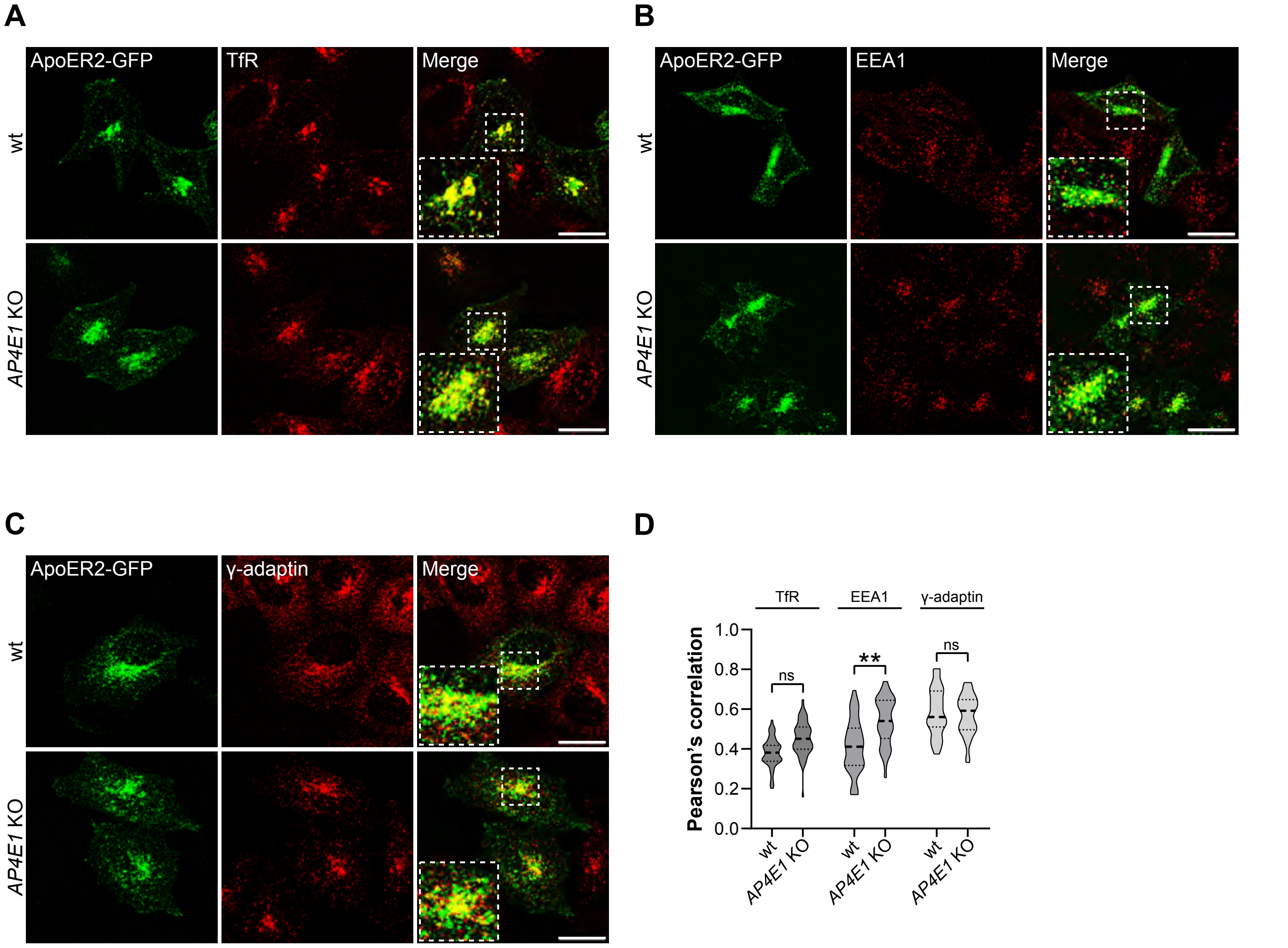

**Figure S2. Increased presence of ApoER2 in Early but not Recycling Endosomes in *AP4E1*-KO HeLa cells.** **(A-C)** HeLa cells were transfected with plasmids encoding wt ApoER2-GFP (green) and immunostained 24 h later with **(A)** TfR as a marker of Recycling endosomes, **(B)** EEA1, as a marker of Early endosomes, and **(C)** the AP1 adaptor subunit γ–adaptin (Red). Scale bar 20 μm. Magnified views of boxed areas are shown in the lower left corner of merged images. **(D)** Graphs showing the Pearson's correlation coefficient obtained from 29-43 cells pooled from n=3 independent experiments. Statistical significance was calculated by Mann-Whitney t-test **p<0.01; ns not significant.

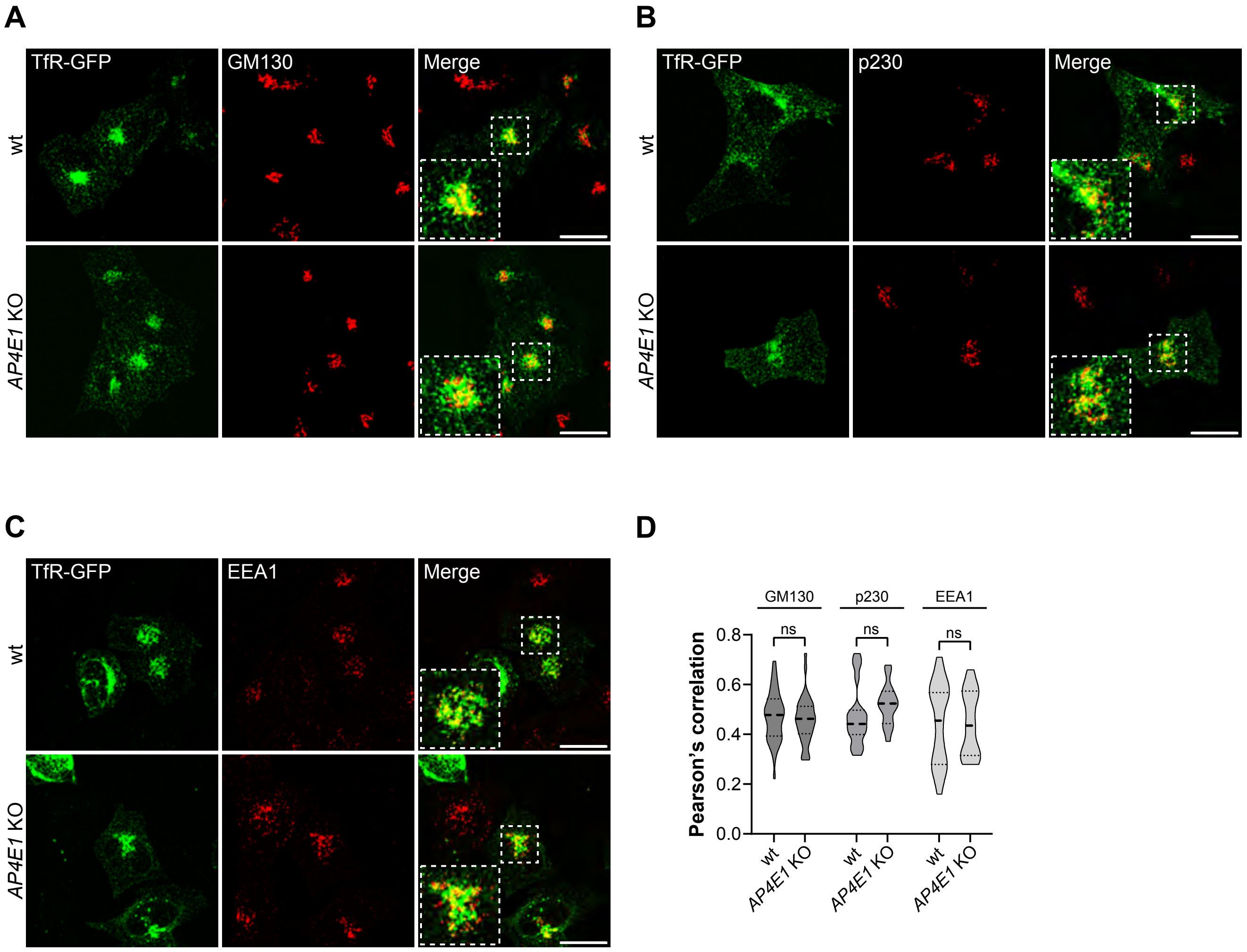

**Figure S3. TfR-GFP does not change its Golgi or Early endosome co-localization in *AP4E*1-KO HeLa cells.** **(A-C)** HeLa cells were transfected with plasmids encoding TfR-GFP (green) and immunostained 24 h later with **(A)** GM130, **(B)** p230, and **(C)** EEA1 (Red). Scale bar 20 µm. Magnified views of boxed areas are shown in the lower left corner of merged images. **(D)** Graphs showing the Pearson's correlation coefficient obtained from 15 to 35 cells pooled from n=2-3 independent experiments. Statistical significance was calculated by Mann-Whitney t-test, ns not significant.

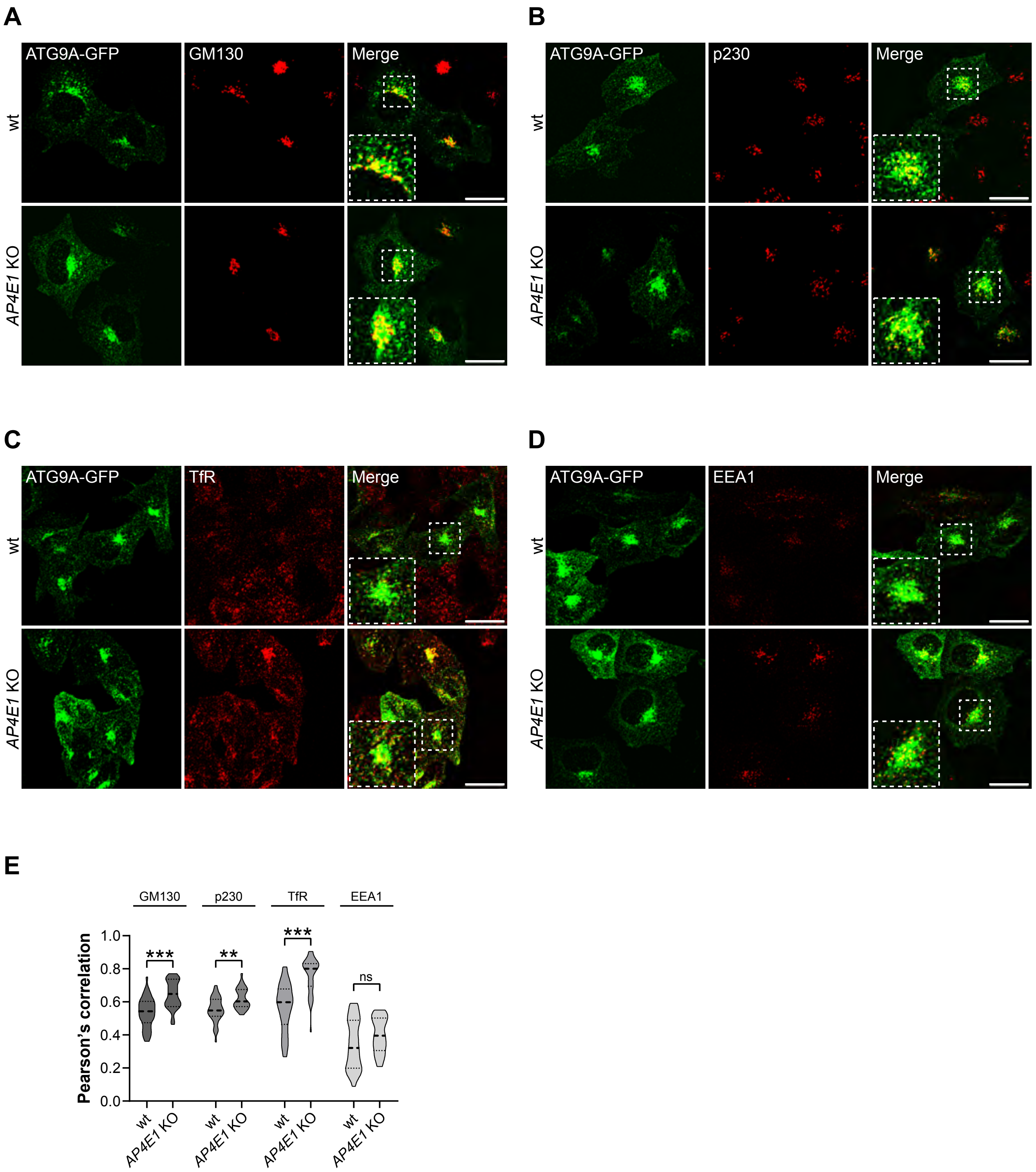

**Figure S4. ATG9-GFP increases its presence in Golgi and recycling endosome in *AP4E1*-KO HeLa cells.** **(A-D)** HeLa cells were transfected with plasmids encoding ATG9A-GFP (green) and immunostained 24 h later with the Golgi markers **(A)** GM130 and **(B)** p230**;** As recycling and early endosomal markers **(C)** TfR and **(D)** EEA1 respectively (Red). Scale bar 20 µm. Magnified views of boxed area are shown in the lower left corner of merged images. (E) Graphs showing the Pearson's correlation coefficient obtained from 30 to 44 cells pooled from n=3 independent experiments. Statistical significance was calculated by Mann-Whitney t-test, **p<0.01, ***p<0.001, **p<0.01, ns not significant.

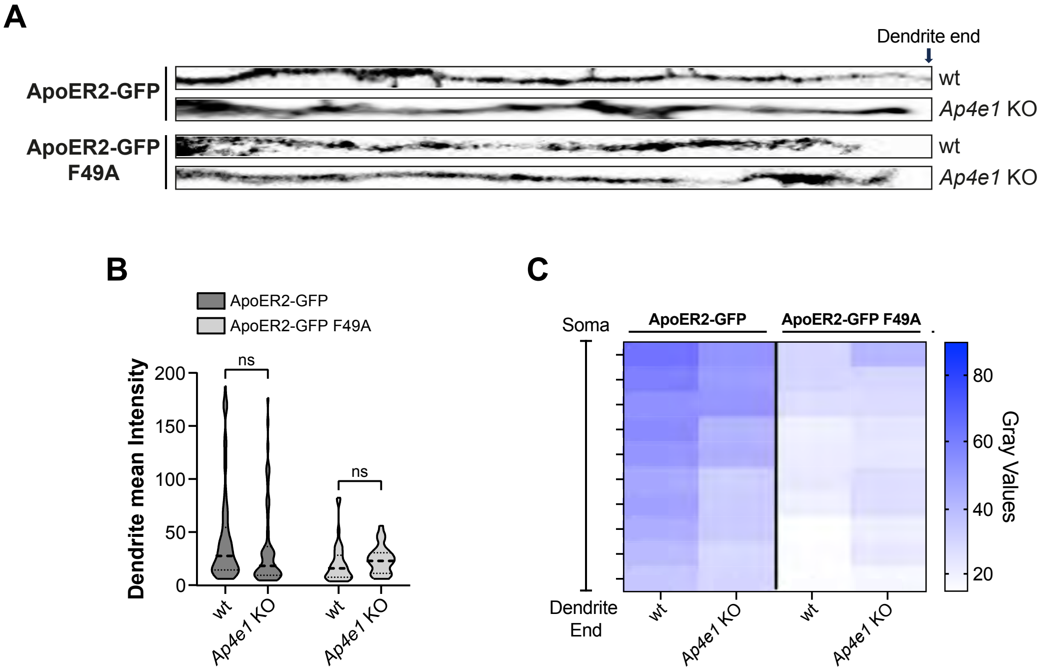

**Figure S5. ApoER2-GFP Dendritic localization is unaffected in *Ap4e1* KO neurons. (A)** wt and *Ap4e1*-KO mouse neurons were transfected at DIV 4 with plasmids encoding HA-ApoER2-GFP or HA-ApoER2-GFP F49A and fixed 24 h later. Representative dendritic segments obtained from neurons in Fig.5. **(B)** Violin Plots showing total intensity measurement of dendritic segments. Averages obtained from 30-75 dendrites and significance was calculated by Mann-Whitney T-test, ns not significant. **(C)** Dendritic GFP signal throughout full length dendrites were binned, averaged and plotted into a heat map to show the spatial distribution of ApoER2 relative to the neuronal soma (see Methods).

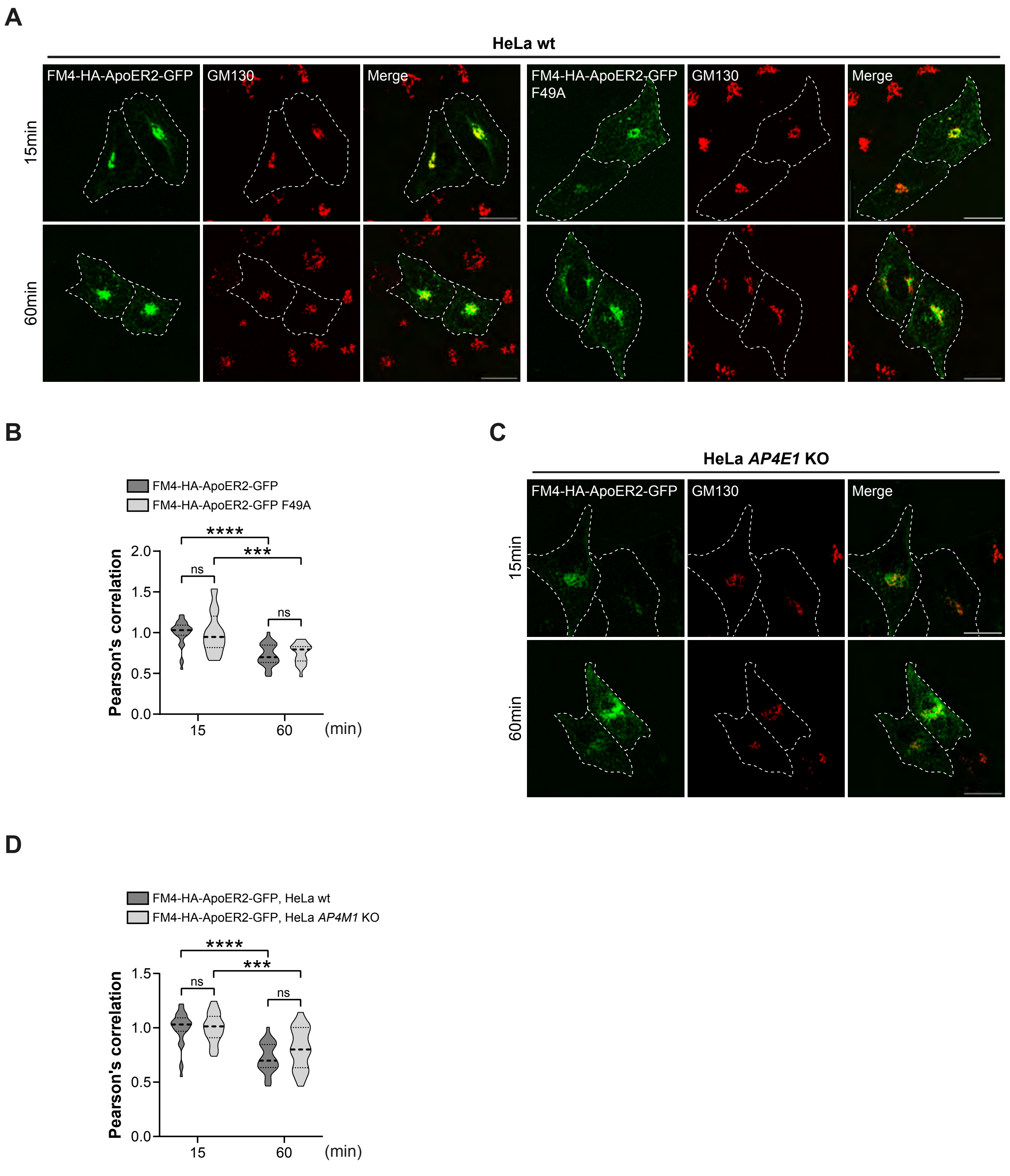

**Figure S6. FM4-HA-ApoER2-GFP TGN exit is not affected in *AP4E1*-KO HeLa cells or by F49A mutation.** **(A)** HeLa wt cells transfected with plasmids FM4-HA-ApoER2-GFP wt or F49A (green) were fixed at the indicated times after addition of DD-Solubilizer, and stained with anti-GM130 (red). Dashed white lines delineate the plasma membrane. Scale bar: 20 µm. **(B)** Pearson's correlation coefficient analysis from 20-34 cells pooled from n=3 independent experiments for each time point and cell line. Statistical significance was calculated by Mann-Whitney t-test , ***$p<0.001$ , ****$p<0.0001$, ns not significant. **(C)** HeLa *AP4E1*-KO transfected with a plasmid encoding FM4-HA-ApoER2-GFP were fixed at the indicated times after addition of DD-Solubilizer. Cells were stained with anti-GM130(red). Dashed white lines delineate the plasma membrane. Scale bar: 20 µm. **(D)** Pearson's correlation coefficient analysis from 25-34 cells pooled from n=3 independent experiments for each time point and cell line. Statistical significance was calculated by Mann-Whitney t-test , ***$p<0.001$, ****$p<0.0001$, ns not significant.

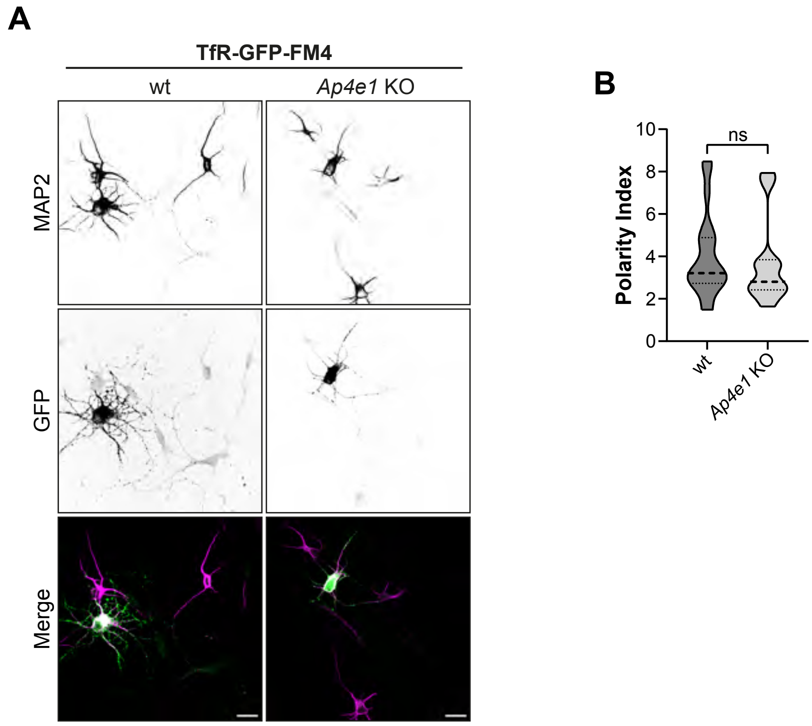

**Figure S7. TfR-GFP-FM4 early secretory trafficking is unaffected in *Ap4e1* KO neurons.** **(A)** TfR-GFP-FM4 was expressed in DIV4 wt and *Ap4e1* KO hippocampal neurons. After 16 h neurons were treated with DD Solubilizer for 2h. Neurons were stained for Scale bar: 20 µm. **(B)** The dendrite/axon polarity index was calculated as described in Methods from 11-19 neurons from 2 independent experiments. Statistical significance was calculated by Mann-Whitney T-test, ns, not significant.

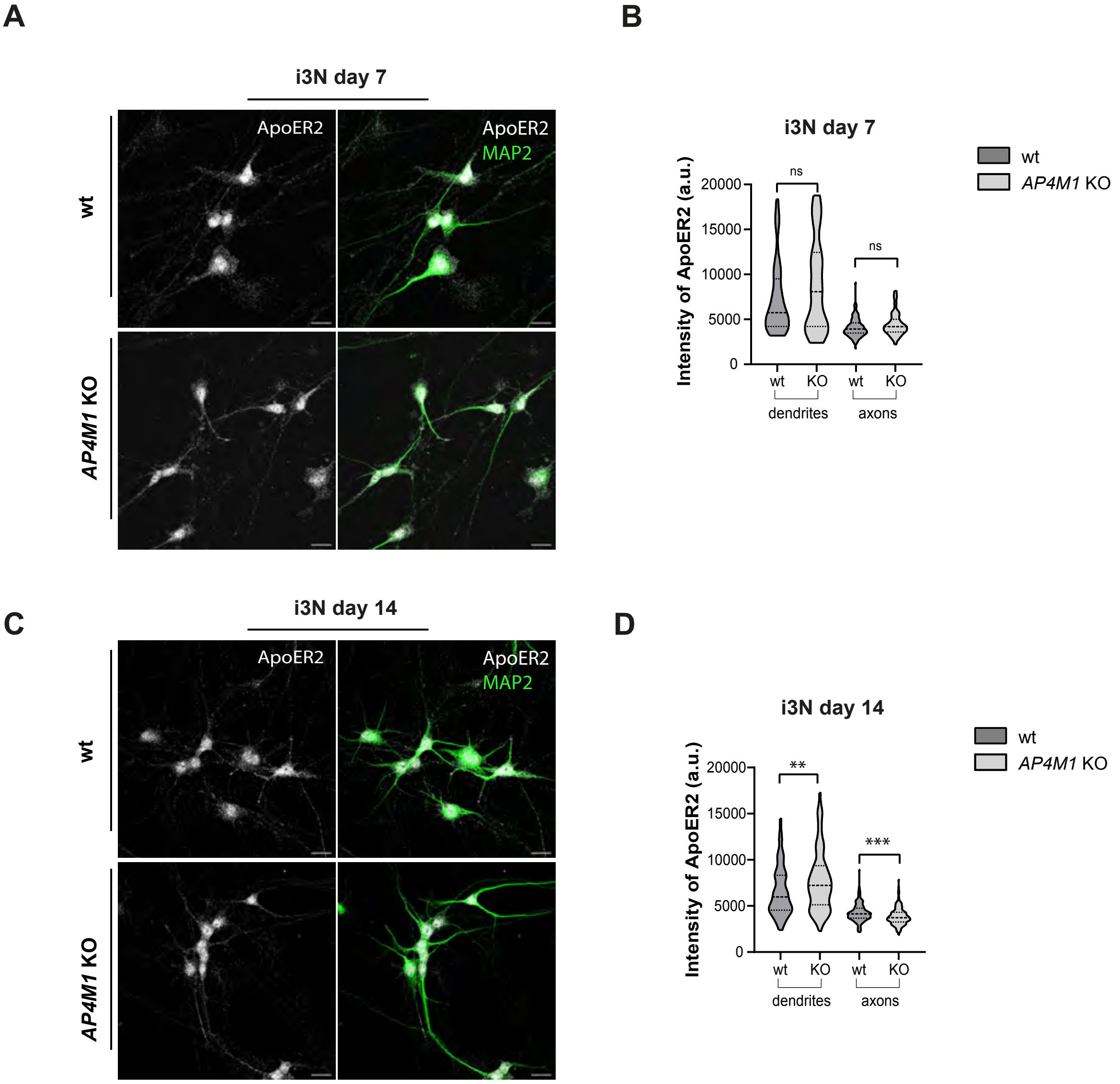

**Figure S8. Dendritic and axonal ApoER2 levels through differentiation of i3Neurons (Related to Fig.9 I,J).** Wild-type and *AP4M1*-KO i3Neurons were differentiated for 7 and 14 days, then stained with anti-ApoER2 (gray) and anti-MAP2 (green). Confocal images were analyzed with Fiji. Intensity of ApoER2 fluorescence was measured in several regions MAP2 positive (dendrites) and MAP2 negative (axons). **(A, B**) After 7 days of differentiation i3Neurons shows no differences on the levels of ApoER2 between dendrites and axons. **(C, D)** After 14 days of differentiation neurons lacking AP-4 show decreased ApoER2 in axons and increased in dendrites compared with the wt neurons. Two experiments, approximately 100 regions of interest analyzed per phenotype in each zone (dendrites or axons) per day of differentiation. Statistical significance was calculated using the Mann-Whitney t-test. $^{**}p<0.01$, $^{***}p<0.001$

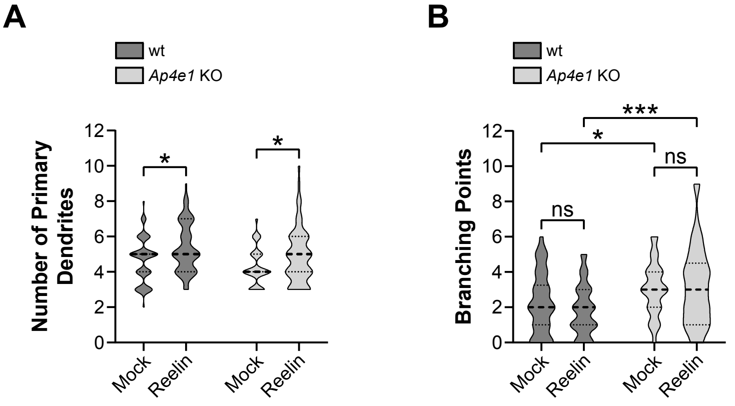

**Figure S9. Analysis of dendritic growth parameters (related to Figure 10 A,B)**. DIV5 hippocampal neurons (from wt and *Ap4e1* KO mice) were incubated with Reelin (10 nM) or mock for 48 h to induce dendritic development. Measurement of **(A)** the number of primary dendrites and **(B)** branching points. Significance was calculated using the ANOVA Tukey multiple comparison test. Results of 50-70 neurons per condition obtained in n=2 independent experiments. $p<0.05$, $^{**}p<0.01$, $^{***}p<0.001$, ns not significant.
